# Perinuclear damage from nuclear envelope deterioration elicits stress responses that contribute to *LMNA* cardiomyopathy

**DOI:** 10.1101/2023.02.14.528563

**Authors:** Kunal Sikder, Elizabeth Phillips, Zhijiu Zhong, Nadan Wang, Jasmine Saunders, David Mothy, Andrew Kossenkov, Timothy Schneider, Zuzana Nichtova, Gyorgy Csordas, Kenneth B. Margulies, Jason C. Choi

## Abstract

Mutations in the *LMNA* gene encoding nuclear lamins A/C cause a diverse array of tissue-selective diseases, with the heart being the most commonly affected organ. Despite progress in understanding the molecular perturbations emanating from *LMNA* mutations, an integrative understanding of the pathogenesis leading to cardiac dysfunction remains elusive. Using a novel cell-type specific *Lmna* deletion mouse model capable of translatome profiling, we found that cardiomyocyte-specific *Lmna* deletion in adult mice led to rapid cardiomyopathy with pathological remodeling. Prior to the onset of cardiac dysfunction, lamin A/C-depleted cardiomyocytes displayed nuclear envelope deterioration, golgi dilation/fragmentation, and CREB3-mediated golgi stress activation. Translatome profiling identified upregulation of Med25, a transcriptional co-factor that can selectively dampen UPR axes. Autophagy is disrupted in the hearts of these mice, which can be recapitulated by disrupting the golgi or inducing nuclear damage by increased matrix stiffness. Systemic administration of pharmacological modulators of autophagy or ER stress significantly improved the cardiac function. These studies support a hypothesis wherein stress responses emanating from the perinuclear space contribute to the development of *LMNA* cardiomyopathy.

**Teaser:** Interplay of stress responses underlying the development of *LMNA* cardiomyopathy

## Introduction

Laminopathy is a collection of human diseases arising from mutations in the *LMNA* gene that encodes A-type lamins A and C (lamin A/C)(*1*). As a major component of the nuclear lamina, a fibrous meshwork lining the inner surface of the nuclear envelope (NE)(*2, 3*), lamin A/C play diverse cellular roles, such as providing protection from mechanical stress(*4–6*) and establishing scaffolding for binding chromatin(*7, 8*) as well as for cell signaling intermediaries(*9–11*). Despite ubiquitous expression in the nuclei of most differentiated mammalian somatic cells, *LMNA* incurs specific mutations that lead to tissue-selective syndromes affecting striated muscle, adipose tissue, and peripheral nerve, as well as multi-system disorders with features of accelerated aging. The most prevalent laminopathy is dilated cardiomyopathy (herein referred to as *LMNA* cardiomyopathy) with variable skeletal muscle involvement. Current therapies are limited to the treatment of congestive heart failure symptoms, hence neither the underlying pathogenesis nor the eventual progression to heart failure are addressed. Targeted therapies based on mechanistic insights that prevent heart muscle deterioration are urgently needed.

Multiple established mouse models of *LMNA* cardiomyopathy that approximate the human disease currently exist. Germline homozygous *Lmna* knockout mice (*Lmna*^-/-^) develop rapid skeletal muscle dystrophy and cardiomyopathy by ∼6 weeks of age(*12, 13*). Germline homozygous knock-in mice with point mutations corresponding to those identified in human patients (i.e. *Lmna*^H222P/H222P^ and *Lmn*a^N195K/N195K^) develop skeletal and cardiac muscle disease with slower kinetics while heterozygotes are unaffected(*14, 15*). An important distinction between the mouse models of *LMNA* cardiomyopathy and the human disease is that in mice, mutations in both alleles of *Lmna* are required for the disease penetrance, whereas in humans the vast majority of *LMNA* cardiomyopathy is autosomal dominant. Despite these differences, mouse models of *LMNA* cardiomyopathy have been instrumental in uncovering a wide array of cellular defects caused by *Lmna* mutations. For example, we and others showed that impaired macroautophagy (autophagy) in the heart underlies *LMNA* cardiomyopathy development in *Lmna*^H222P/H222P^ and *Lmn*a^-/-^ mice(*16, 17*) but why and how this impairment contributes to disease pathogenesis is unknown.

Nuclear envelope fragility/damage resulting from *LMNA* mutations is widely believed to be involved in the disease pathogenesis but how it contributes is not well understood(*18*). Nuclear envelope perturbations from defects in lamin A/C have been noted in cell culture models(*5, 6, 19*) as well as in the myocardium from murine models(*14, 20, 21*) and from humans(*22*), but how these alterations engender pathogenic mechanisms remain unclear. A detailed kinetic study using a temporally controlled, cardiomyocyte (CM)-specific deletion model of *Lmna* would enable accurate dissection of molecular pathogenesis emanating from nuclear fragility. To this end, we developed a novel cre recombinase driver line referred to as CM-CreTRAP mice. It is a bi-cistronic transgenic line in which tamoxifen (Tam)-regulatable Cre recombinase (CreERT2) and EGFP-L10a fusion protein are co-expressed under the control of the myosin heavy chain 6 (*Myh6*) promoter thereby directing their expression specifically in CMs. EGFP-L10a is enhanced green fluorescent protein fused to ribosomal protein L10a, a component of the 60S ribosomal protein, and the CM-specific driven expression of this fusion protein allows tagging of polysomes for immunoaffinity purification of translating mRNA (<u>T</u>ranslating <u>R</u>ibosome <u>A</u>ffinity <u>P</u>urification [TRAP])(*23*). Initially developed by the laboratory of Dr. Nathaniel Heintz for the purpose of characterizing neuronal subpopulations in mice(*24, 25*), the technique has been adapted to other model organisms including *Drosophila melanogaster*, *Xenopus laevis*, *Danio rerio*, and *Arabidopsis thaliana*(*26–30*).

Additionally, the method has evolved to enable Cre-directed EGFP-L10a tagging(*31*) and tetracycline operator-mediated temporal regulation(*32*). Following intercross with the *Lmna*^flox/flox^ line, we induced CM-specific *Lmna* deletion in 12-week-old adult mice and detailed the development of *LMNA* cardiomyopathy. Using a combination of *in vivo* animal model, freshly isolated primary CMs, and *in vitro* culture models, we show that *Lmna* deletion in adult CMs cause significant damage to the nuclear envelope and the perinuclear space. The golgi apparatus is particularly affected, leading to a CREB3-mediated golgi-stress response. Furthermore, *in vitro* disruption of golgi recapitulates the disrupted autophagy phenotype observed in the germline mouse models of *LMNA* cardiomyopathy. Pharmacological modulators of autophagy or ER stress significantly improved the cardiac function, suggesting these pathways contribute to the pathogenesis and provide an additional avenue of therapeutic approaches.

## Results

### Generation of a murine model with CM-specific Lmna deletion

To gain mechanistic insights into *LMNA* cardiomyopathy in a cell type-specific fashion, we engineered CM-CreTRAP mice that express a bi-cistronic construct consisting of CreERT2 and EGFP-L10a linked by an IRES (Supplementary Fig. 1a). The transgene’s expression is driven by the *Myh6* promoter(*33*) and stabilized by the human growth hormone polyadenylation signal. We identified 2 founder lines (line 1 and line 2) with stable transgene transmission and robust expression of EGFP and CreERT2 in line 1 and lesser expression of these proteins in line 2 (Supplementary Fig. 1b). Direct fluorescence images of frozen heart sections confirmed the transgene protein expression in both founder lines (Supplementary Fig. 1c). The transgene mRNA expression in line 1 using primers that recognize EGFP-L10a sequences was specific to the heart with undetectable expression in other tissues tested (Supplementary Fig. 1d). Immunofluorescence staining on frozen heart sections from heterozygous transgenic positive CM-CreTRAP mice revealed a robust GFP signal that colocalized with sarcomeric actin, indicating CM-specific transgene expression (Supplementary Fig. 1e). We also noticed punctate signals within the nucleus (inset, white arrows), which is consistent with the established pattern of EGFP-L10a expression(*25, 31*).

### Rapid myocardial dysfunction with robust fibrosis following Lmna deletion in adult CMs

We assessed the impact of *Lmna* deletion in CMs of CM-CreTRAP mice interbred with *Lmna*^flox/flox^ mice(*34*). All mice with Cre driving transgenic allele (CM-CreTRAP) were studied as heterozygotes. 100 mg/kg/day Tam was delivered intraperitoneally for five consecutive days followed by a 2 day rest (designate as “0 week”) in 12-weeks-old line 1 CM-CreTRAP:*Lmna*^flox/+^, CM-CreTRAP:*Lmna*^flox/flox^, mice and corn-oil for vehicle treated CM-CreTRAP:*Lmna*^flox/flox^ mice (Fig. 1a). To assess *Lmna* deletion efficiency, we isolated CMs from a subset of Tam and vehicle-treated CM-CreTRAP:*Lmna*^flox/flox^ mice for immunoblot analysis and observed ∼86% and ∼78% reduction in line 1 and line 2, respectively (Supplementary Fig. 2a). From this point on, most of our analyses were performed in line 1. With the confirmation of lamin A/C depletion in CMs, we then assessed its impact on the myocardium. The heart size was not significantly different at 2 weeks post Tam but by 4 weeks, the enlargement was obvious (Fig. 1b). This enlargement was confirmed by heart weight/tibia length and heart weight/body weight ratio measurements (Supplementary Fig. 2b).

**Fig. 1.**
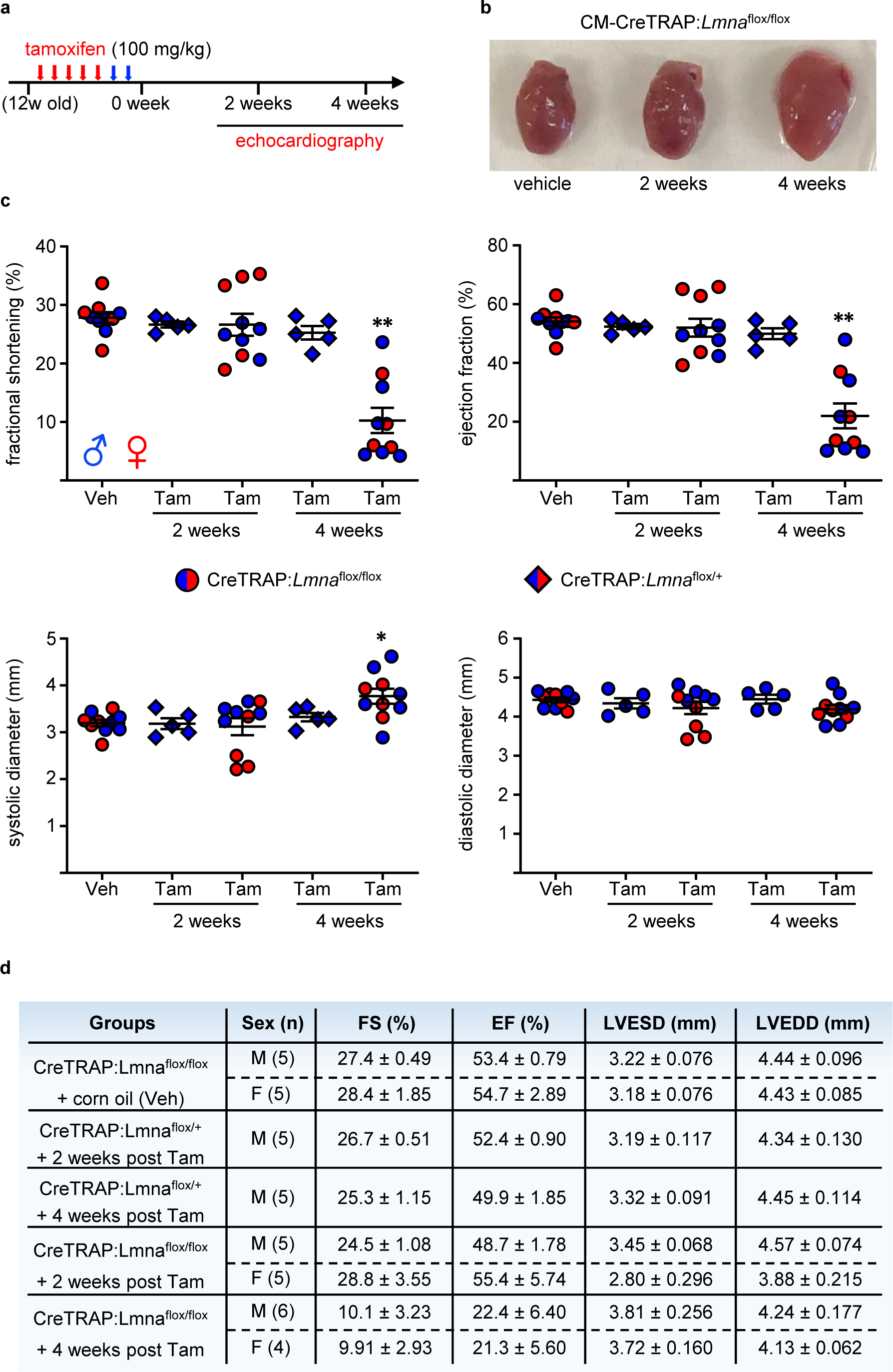
Rapid cardiac performance deterioration after CM-specific *Lmna* deletion. (**a**) Tam dosing schedule. Red arrows denote days of Tam injection and blue arrows show 2 days of rest. “12w old” denotes 12-week-old mice. (**b**) Hearts excised from CM- CreTRAP:*Lmna*^flox/flox^ mice treated with vehicle (Veh) or Tam for 2 and 4 weeks. (**c**) Echo on CM-CreTRAP:*Lmna*^flox/+^ (diamond) and CM-CreTRAP:*Lmna*^flox/flox^ (circle) at 2 and 4 weeks Tam treatment. * and ** denotes p = 0.015 and p < 0.0001, respectively, using one way ANOVA followed by Dunnett multiple comparison correction with the Veh group serving as comparison control. Error bars = SEM. Blue = male and red = female mice. (**d**) Tabulation of echo data ± SEM. FS = fractional shortening; EF = ejection fraction; LVESD = left ventricular end systolic dimension; LVEDD = left ventricular end diastolic dimension.

We then measured cardiac function by performing M-mode echocardiography (echo) at these timepoints to calculate fractional shortening (% reduction of left ventricular dimension at systole), ejection fraction (estimated calculation of % of blood ejected from left ventricle), and the systolic/diastolic left ventricular dimensions required to derive the first two parameters (Fig. 1c). Tam-treated CM-CreTRAP:*Lmna*^flox/flox^ mice revealed consistent and reproducible kinetics of cardiac dysfunction with a rapid decline in cardiac function by 4 weeks as determined by fractional shortening and ejection fraction (Figs. 1c, d, and Supplementary Table 1). Moreover, despite prior reports of Tam and Cre toxicity in the heart(*35, 36*), vehicle only-treated CM-CreTRAP:*Lmna*^flox/flox^ as well as Tam-treated CM-CreTRAP:*Lmna*^flox/+^ mice revealed no cardiac dysfunction (Figs. 1c, d), indicating that only the homozygous loss of *Lmna* in CMs caused rapid and severe cardiomyopathy in our experimental conditions.

To correlate organ function with tissue morphology, we performed histopathology assessing tissue structure, collagen deposition, and hypertrophy using Hematoxylin and Eosin (H&E), Picro-Sirius Red /Masson’s trichrome, and wheat germ agglutinin (WGA) staining, respectively (Fig. 2a). H&E revealed obvious abnormalities at 2 weeks post Tam in the form of myocyte degeneration (blue arrows) and the presence of pyknotic nuclei (yellow arrows) (Fig. 2a). By 4 weeks, significant myocyte swelling and loss were obvious with abundant cytoplasmic vacuolation (Fig. 2a, green arrows). These results indicate that histological abnormalities in the myocardium are present at 2 weeks post Tam treatment despite the lack of cardiac functional changes. Furthermore, membrane staining with WGA revealed a progressive increase in the myocyte cross-sectional area, demonstrating myocyte swelling/hypertrophy (Fig. 2a). Picro-Sirius Red staining revealed a subtle increase in collagen deposition in the interstitial space as early at 1 week post Tam that progressively increased, with substantial fibrosis by 4 weeks (Fig. 2a). Masson’s trichrome stain of the same hearts revealed a similar pattern (Supplementary Fig. 3a) with quantitation determined to be ∼28% fibrosis (Fig. 2b). Furthermore, measurement of the myocyte cross-sectional area revealed a significant increase at 2 weeks post Tam (Fig. 2c), further demonstrating that, although cardiac function was normal at this timepoint, histological abnormalities have manifested. Altogether, these observations indicate a progressive pathological remodeling in response to CM-specific lamin A/C depletion. No abnormalities were observed in quadriceps from these mice, demonstrating specificity of the disease in the heart (Supplementary Fig. 3b). Moreover, hearts of Tam-treated CM- CreTRAP:*Lmna*^flox/+^ mice displayed no obvious histological abnormalities (Supplementary Fig. 3c), further confirming that the myocardial pathogenesis requires the loss of both *Lmna* alleles and that Tam as well as the CreERT2 expression by themselves had no observable effect on the heart. Similar fibrotic responses were also observed in transgenic line 2 (Supplementary Fig. 3d).

**Fig. 2.**
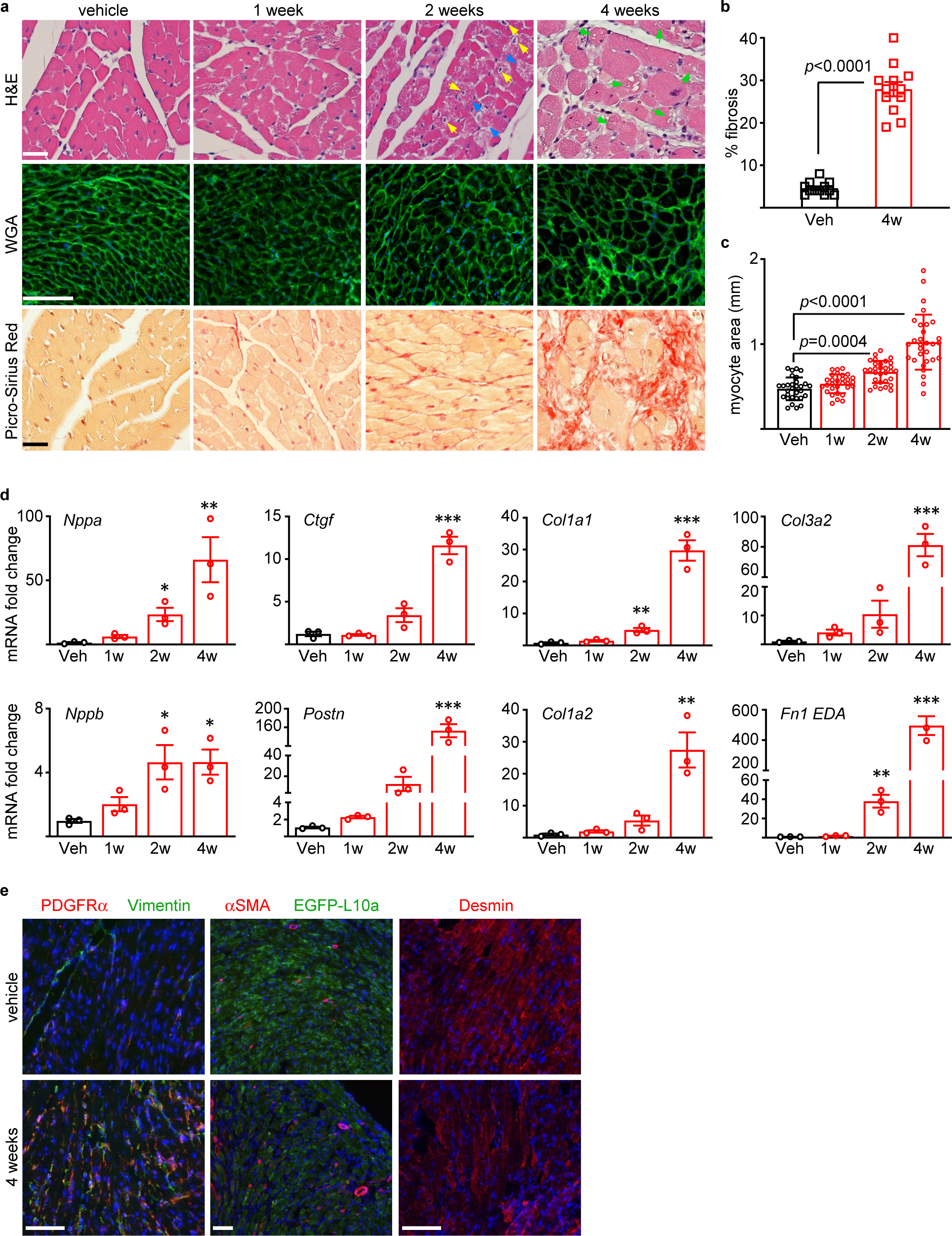
Pathological cardiac remodeling accompanies cardiac dysfunction. (**a**) Histological analyses of the hearts from CM-CreTRAP:*Lmna*^flox/flox^ mice at 1, 2, and 4 weeks post Tam or vehicle treatment. Blue, yellow, and green arrows denote degenerating myocytes, pyknotic nuclei, and vacuolation, respectively. H&E = Hematoxylin and Eosin; WGA = wheat germ agglutinin. Scale bar = 50 μm. Representative images are shown. (**b**) Quantitation of average % fibrosis of Masson’s Trichrome staining from 12 independent images from 3 mouse hearts per group. Error bars = SD. % Fibrosis *p* value was determined using unpaired, two-tailed Student’s t test. (**c**) Quantitation of myocyte cross- sectional area based on WGA staining. 3 images were taken per section and average area of 30 individual CMs were measured in total per group. Error bars = SD. *p* values were determined using one way ANOVA and Dunnett correction with Veh as control. (**d**) qPCR of cardiac stress and profibrotic markers in hearts of Tam-treated CM- CreTRAP:*Lmna*^flox/flox^ mice. 1w-4w denote weeks post final Tam dosing. Data are presented as fold change relative to Veh. *, **, and *** denote p < 0.05, 0.01, and p < 0.0001, respectively, using one way ANOVA with Dunnett correction with Veh as control. Error bars = SEM. n = 3. (**e**) Immunofluorescence images of PDGFRα, vimentin, α-smooth muscle actin (αSMA), EGFP-L10a, and desmin on mouse hearts 4 weeks after vehicle or Tam treatment. Scale bar = 100μm.

The above histological changes were correlated with increased transcript expression of cardiac stress markers as well as markers of pathological fibrosis (Fig. 2d). Natriuretic peptide precursors *Nppa* and *Nppb*, which are produced in response to excessive CM stretch, were enhanced by 2 weeks post Tam treatment and increased further by 4 weeks. A similar pattern was observed for markers of activated cardiac fibroblasts (*Postn* and *Ctgf*) and extracellular matrix synthesis (*Col1a1*, and *Col1a2*) as well as those associated with pathological remodeling (*Col3a2* and *Fn1*-EDA). These changes were absent in Tam-treated CM-CreTRAP:*Lmna*^flox/+^ mice, demonstrating that again, in our system, Tam treatment or CreERT2 expression did not elicit pathological remodeling (Supplementary Fig. 3e). Indirect immunofluorescence revealed increased α- smooth muscle actin (myofibroblast marker), vimentin (mesenchymal marker expressed in cardiac fibroblasts), and PDGFRα (also a cardiac fibroblast-selective marker) staining in the myocardial sections from CM-CreTRAP:*Lmna*^flox/flox^ mice at 4 weeks post Tam dosing, indicating increased presence of cardiac fibroblasts and myofibroblasts (Fig. 2e). Desmin staining further revealed disruption of typical CM alignment and distortions of CM shape owing to its degeneration (Fig. 2e).

### NE abnormalities and partial UPR activation in Lmna-deleted CMs

A closer inspection of the higher magnification H&E-stained hearts revealed nuclear abnormalities uncharacteristic of those previously associated with *LMNA* cardiomyopathy models. Previous studies using germline knockout and knockin mouse models revealed abnormal elongation of nuclei in CMs(*21, 37*). In the hearts of Tam- treated CreTRAP:*Lmna*^flox/flox^ mice, a greater variety of nuclear shape abnormalities at higher frequencies were noted (Fig. 3a). At 2 weeks post Tam, nuclear shapes were much more irregular, with many being swollen, pyknotic, or grossly misshapen (Fig. 3a). Even accounting for sectioning artefacts, abnormal nuclei were more prevalent in the Tam- treated hearts than those treated with corn-oil. The severity and the frequency of these abnormalities became more pronounced at 4 weeks post Tam treatment (Supplementary Fig. 4a).

**Fig. 3.**
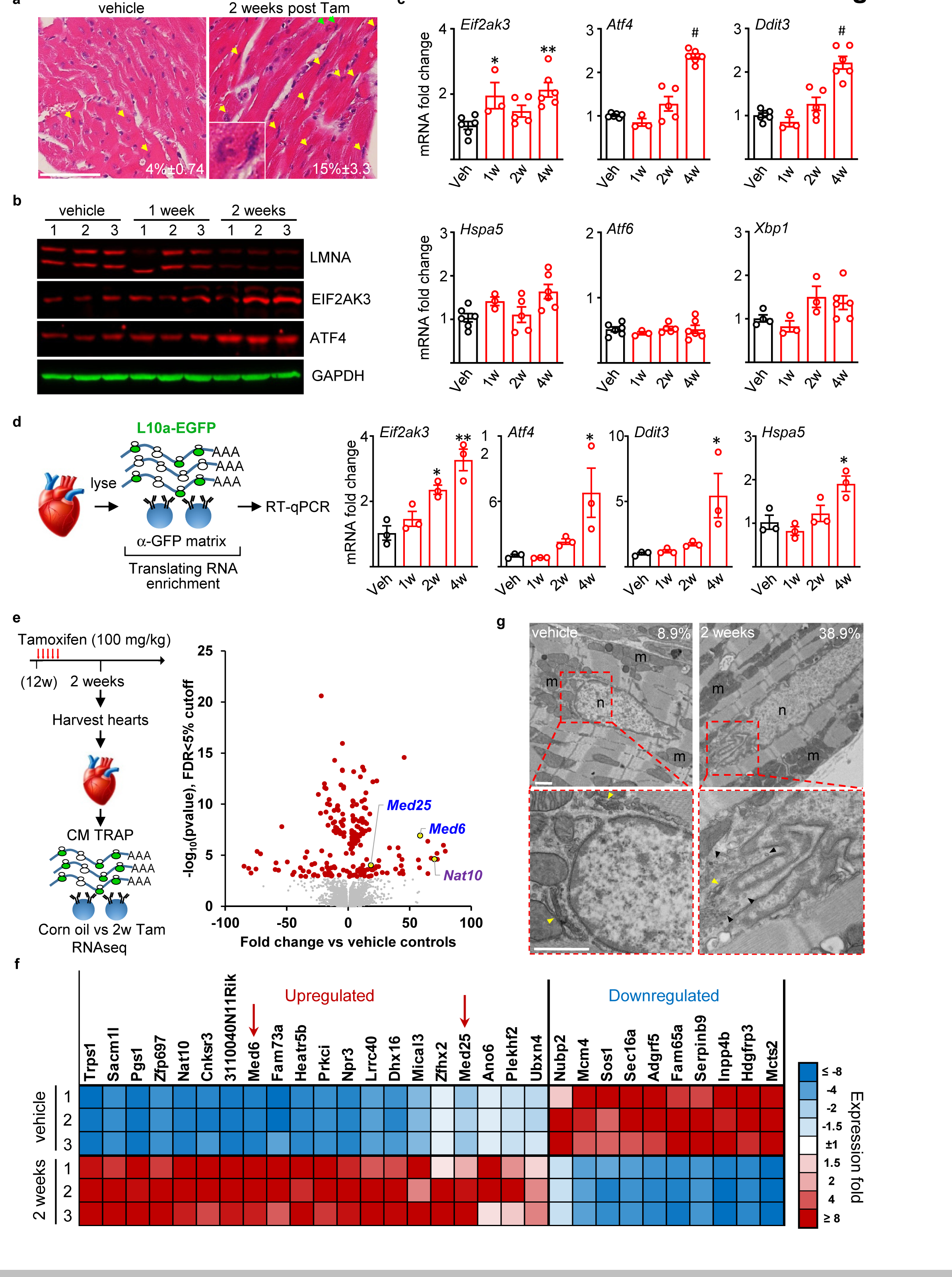
Selective activation of UPR in laminopathic hearts. (**a**) H&E staining of longitudinal sections of hearts from CM-CreTRAP: *Lmna*^flox/flox^ mice 2 weeks post vehicle or Tam treatment. Yellow arrows highlight abnormally shaped, ruptured, and stretch nuclei with green arrows showing nuclei in inset. Average number ± SEM of abnormal nuclei per field counted from at least 15 fields from 3 mice per group are shown within the images. (**b**) Immunoblot of LMNA (lamin A/C), EIF2AK3 (PERK), and ATF4 with GAPDH as internal control in hearts from vehicle or Tam-treated CM-CreTRAP: *Lmna*^flox/flox^ mice. Numbers on top of blots denote individual heart samples. 1 and 2 weeks denote time post final Tam treatment. (**c**) qPCR analyses of transcripts encoding mediators of unfolded protein response in hearts from CM-CreTRAP: *Lmna*^flox/flox^ mice treated with vehicle (Veh) or Tam. 1w, 2w, and 4w denote weeks post Tam treatment. *, **, and # denote p values of < 0.05, 0.005, and 0.0001, respectively, using one way ANOVA with Dunnett’s post hoc with Veh as control. n=5 except for 1w (n=3). All error bars in this figure = SEM. (**d**) Schema of experimental design using TRAP. Right panel shows qPCR analyses on CM-specific translating mRNAs probed for *Eif2ak3*, *Atf4*, *Ddit3*, and *Hspa5* encoding PERK, ATF4, CHOP, and BiP, respectively, from CM-CreTRAP: *Lmna*^flox/flox^ mice 1 - 4 weeks post Tam treatment. * and ** denote p < 0.05 and p < 0.0005, respectively, using one way ANOVA with Dunnett’s post hoc with Veh as control. n=3. (**e**) Left panel - experimental schema of TRAP/RNAseq and right panel - volcano plot of differentially expressed genes sorted by –log_10_ of nominal p value with 5% false discovery rate (FDR) cut off. (**f**) Heatmap of top 30 coding genes filtered by adjusted p values of perfect markers. Red arrows denote the position of *Med6* and *Med25*. (**g**) TEM micrographs of hearts from CM-CreTRAP: *Lmna*^flox/flox^ mice 2 weeks post vehicle or Tam treatment. Perinuclear regions denoted by dashed red boxes are shown on the bottom. Yellow and black arrowheads denote the golgi and small vesicles, respectively. m = mitochondria, n = nucleus. Scale bar = 1 μm. The numbers in line with the EM micrographs denote percentage of abnormal/damaged nuclei quantified in vehicle (n = 45 total nuclei) and 2 weeks post Tam (n = 31 total nuclei) treatment groups 2 independent hearts per group.

The nuclear membranes are contiguous with the rough endoplasmic reticulum (ER), representing distinct regions of a single membrane system(*38*). We reasoned that the nuclear shape abnormalities and the damage arising from them would also cause ER-nuclear membrane disruption leading to ER stress. Moreover, evidence of ER stress activation has been documented in hearts of transgenic models of *LMNA* cardiomyopathy(*20*) as well as in patient-derived fibroblasts(*39*). We therefore assessed for ER stress activation in hearts of CreTRAP:*Lmna*^flox/flox^ mice following *Lmn*a deletion. ER stress triggers a set of signaling pathways, collectively termed unfolded protein response (UPR), for which there are three canonical branches; PERK, ATF6, and IRE1(*40*). Following CM-specific *Lmna* deletion, we observed elevated protein expression of PERK (EIR2AK3) and its downstream mediator ATF4 at 2 weeks post Tam dosing (Fig. 3b). qPCR analyses further indicated that *Eif2ak3*, *Atf4*, and *Ddit3* (encoding CHOP, a downstream target of ATF4) expression were elevated at the transcript level (Fig. 3c).

To assess the ATF6 branch of the UPR, we measured mRNA levels of genes induced by ATF6. Transcriptionally active ATF6 can transactivate *Xbp1*, *Hspa5* (encoding BiP), as well as its own transcription to establish a positive feedback loop(*41–43*). Expression analysis revealed little changes in the mRNA levels of *Xbp1*, *Hspa5*, and *Atf6* in the CM-specific *Lmna*-deleted hearts (Fig. 3c), suggesting that the ATF6 branch is not robustly activated. To probe the IRE1 branch, we measured *Xbp1* splicing as a proxy for IRE1 activation(*44*) by employing PCR primers that span the intron excised by activated IRE1(*45*). Although *Xbp1* splicing was readily detectable in brefeldin A (BFA)-treated wildtype neonatal CMs (nCMs), this splicing was dampened in lamin A/C-depleted nCMs (Supplementary Fig. 4b) or absent in hearts from Tam-treated CreTRAP:*Lmna*^flox/flox^ mice (Supplementary Fig. 4c). Hearts from Tam-treated CM-CreTRAP:*Lmna*^flox/+^ mice did not display increased expression of any of these ER stress markers, demonstrating that Tam or CreERT2, in and of themselves, did not trigger ER stress (Supplementary Fig. 4d). Our collective results suggest that CM-specific *Lmna* deletion elicits selective UPR activation towards the PERK axis. We observed similar findings in the human disease, as the protein levels of EIF2AK3 and ATF4 in hearts from patients with *LMNA* cardiomyopathy were elevated (∼1.6 fold and ∼2.2 fold for EIF2AK3 and ATF4, respectively) relative to age- and sex-matched healthy controls (Supplementary Fig. 4e), supporting our observations in the mouse model.

To remove the influence of non-myocytes in our analyses, we isolated CM-specific translating mRNA using TRAP from vehicle and Tam-treated mice and performed RT-qPCR (Fig. 3d). To validate the immunopurified mRNA are CM-specific, we interrogated TRAP mRNA with primers specific to markers of various cell types in the myocardium. The TRAP mRNAs were enriched with those encoding CM-specific markers with little to no cardiac fibroblast or endothelial cell-specific markers (Supplementary Fig. 4f), confirming that the isolated mRNA are specifically from CMs. From 2 weeks post Tam treatment and onwards, we observed increases in *Eif2ak3*, *Atf4*, *Ddit3*, and to a lesser degree *Hspa5* expression (Fig. 3d), further supporting that some form of UPR is activated in *Lmna*-deleted CMs.

Given the selective UPR activation signature specifically in CMs, we sought to obtain a broader snapshot of the gene expression profile. To achieve this, we performed RNAseq following TRAP. Translating mRNA isolated from ventricular tissue of mice 2 weeks after treatment with vehicle or Tam were profiled and compared (Fig. 3e). This timepoint was chosen based on our results showing lamin A/C depletion in CMs but prior to detectable deterioration in cardiac function. The goal was to capture early perturbations elicited by lamin A/C depletion before large changes are brought on by cardiac functional decline. Data for sequencing depth, alignment rates, and complexity curves for the samples as well as PCA analysis demonstrate highly concordant behavior in replicates (Supplementary Fig. 5a, b). However, we did notice large fold changes for some genes and a gap in the FDR distribution, which we surmise stem from lower complexity reads (Supplementary Fig. 5c). Raw sequencing results (Supplementary File 1) show that a large proportion of the reads are dominated by mRNAs encoding sarcomeric, mitochondrial, and CM-specific isoform/enriched proteins (Supplementary Fig. 5d). This is expected from a CM-specific translatome and demonstrates that our TRAP worked as intended, which was to purify CM-specific translating mRNA from whole tissue. Nevertheless, to further ensure against false positives, we stringently filtered the genes by perfect markers, which is defined as all experimental replicates displaying same trends in expression over controls.

A volcano plot shows a pair of genes, *Med6* and *Med25*, which were significantly upregulated by 2 weeks (Fig. 3e, right panel). We also noted increased expression of *Nat10*, which was previously linked to misshapen nuclei in lamin A/C-depleted cells(*46*). A heat map of top perfect markers showed that *Med6* and *Med25* are ranked 8^th^ and 17^th^, respectively, as sorted by false discovery rate (FDR)-adjusted p values for upregulated genes (Fig. 3f, red arrows). We confirmed their protein expression in hearts of both Tam- treated CM-CreTRAP:*Lmna*^flox/flox^ mouse lines 1 and line 2 (Supplementary Fig. 6). MED25 and MED6 are members of the Mediator complex, an evolutionarily conserved transcriptional co-factors previously identified as regulators of UPR(*47*), further supporting the presence of UPR activation in the hearts of Tam-treated CM- CreTRAP:*Lmna*^flox/flox^ mice.

In light of the selective UPR activation observed in CMs, we sought to obtain a more detailed view of the nuclear damage, particularly on the perinuclear space, by using transmission electron microscopy (TEM) (Fig. 3g). Similar to H&E, we observed increased NE ruffling/breakdown from 2 weeks post Tam treatment (Fig. 3g and Supplementary Fig. 7a, b), which is prior to detectable cardiac functional decline and obvious pathological remodeling. By 4 weeks post Tam, the frequency of NE undergoing breakdown increased further (Supplementary Fig. 7a and b). Vacuoles of varying size (1 to 3 μm) adjacent to the NE and at the perinuclear space were present. Also present in the perinuclear space was an abundance of small (∼100 nm) vesicles that congest the ER-golgi interface, particularly around areas of NE ruffling/breakdown (Fig. 3g and Supplementary Fig. 7a). These vesicles were abundant at both 2 and 4 weeks post Tam but far less so in vehicle-treated hearts, suggesting that the general abnormalities at the perinuclear region are associated with disease pathogenesis. Altogether, our data show that CM-specific *Lmna* deletion results in NE breakdown, perturbations at the perinuclear space, and selective UPR activation all prior to detectable cardiac functional decline.

### Golgi stress in response to Lmna deletion

In adult CMs, the perinuclear endoplasmic reticulum (ER) and golgi are tightly pack into a confined space around the perinuclear space. The area is also filled with perinuclear mitochondria in a fusiform shape (Supplementary Fig. 8a), which appear to have unique functions relative to their intrafibrillar counterparts(*48*). Given that partial UPR activation in response to *Lmna* deletion in CMs is characteristic of a golgi stress response(*49, 50*), we reasoned that golgi may also be perturbed. We analyzed our EM data with a particular focus on the golgi and observed frequent golgi stack dilation/fragmentation starting from 2 weeks post Tam that increased further in frequency at 4 weeks (Fig. 4a and Supplementary Fig. 8b, c). Furthermore, the dilation of the golgi stacks appeared to be mostly localized around areas adjacent to NE deterioration (Supplementary Fig. 8b, c), indicating that golgi swelling and NE damage may be pathologically linked.

**Fig. 4.**
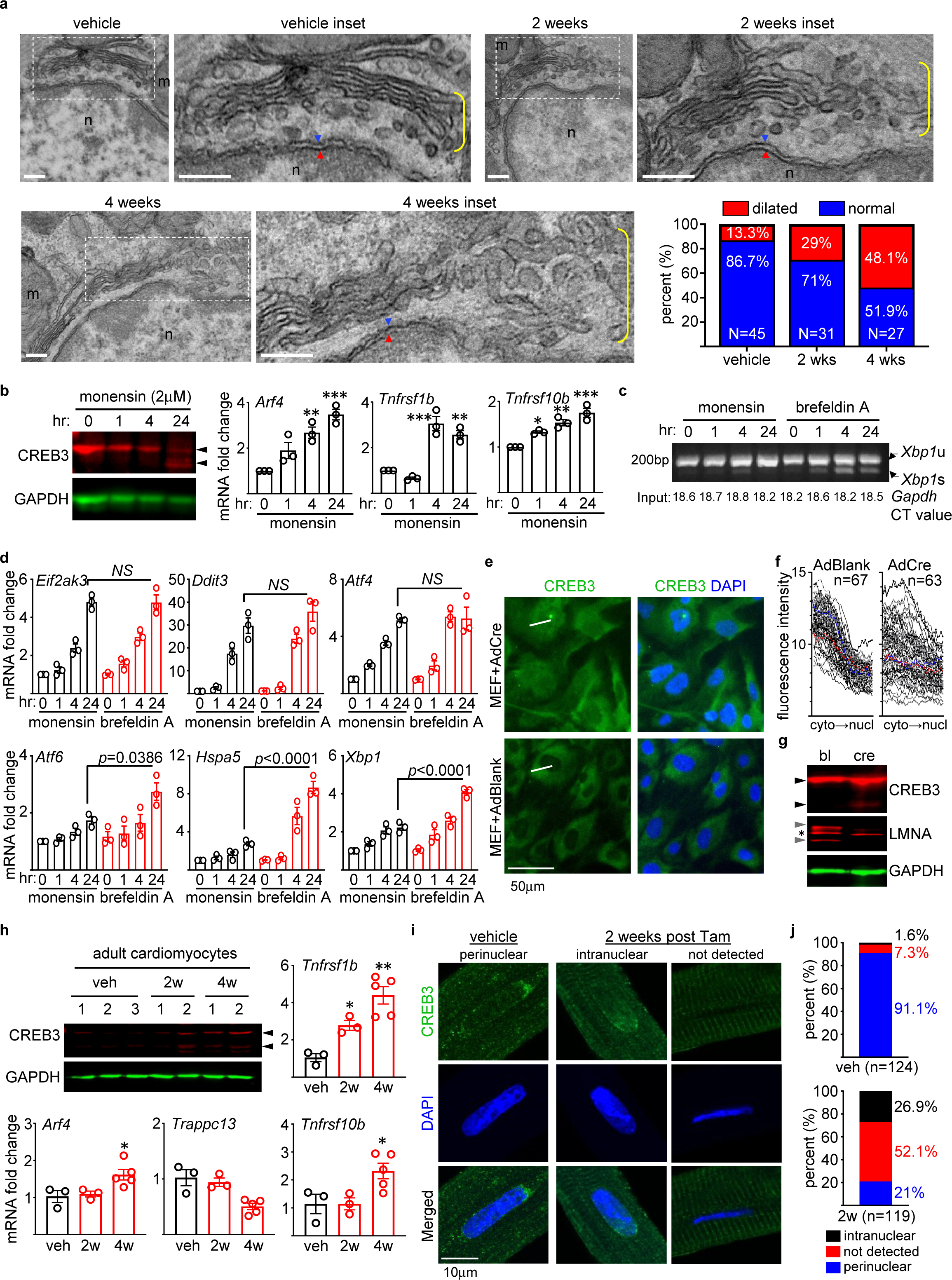
Golgi damage and CREB3 activation in CMs with *Lmna* deletion. (**a**) TEM images of hearts from CM-CreTRAP: *Lmna*^flox/flox^ mice treated with vehicle, 2 and 4 weeks post Tam. Dotted boxes denote insets and yellow brackets denote golgi stacks. Blue and red arrowheads denote outer and inner nuclear membranes, respectively. m = mitochondria, n = nucleus. Scale bars = 200 nm. Quantitation of EM images showing % of dilated and normal golgi apparatus in vehicle (n = 45 total nuclei) as well as 2 weeks (n=31) and 4 weeks (n = 27) post Tam treatment. (**b**) Left panel - immunoblot of CREB3 and GAPDH in *Lmna*^flox/flox^ MEFs treated with 2 μM monensin. Black arrowheads denote CREB3 and its cleavage product. Right panels show qPCR analyses probed for *Arf4*, *Tnfrsf1b*, and *Tnfrsf10b*. *, **, and *** denote p < 0.05, p < 0.005, and p < 0.0005, respectively, using one way ANOVA with Dunnett’s post hoc with 0 hr as the reference control. n=3. (**c**) RT-PCR analyses of *Xbp1* mRNA splicing in *Lmna*^flox/flox^ MEFs treated as described in 4b. *Xbp1*u and *Xbp1*s denote unspliced and spliced variants, respectively. Numbers on the bottom of gel denote CT values for *Gapdh* as input control. Representative images from 3 independent experiments are shown. (**d**) qPCR analyses of transcripts encoding mediators of UPR in *Lmna*^flox/flox^ MEFs treated with either monensin (2 μM) or brefeldin A (20 μM) for 1, 4, and 24 hrs. 0 hr denotes vehicle treatment (EtOH for monensin and DMSO for brefeldin A). The indicated p values were derived using one way ANOVA with Dunnett’s post hoc with 0 hr monensin as reference. n=3. NS = not significant. (**e**) Representative immunofluorescence images of CREB3 and DAPI on *Lmna*^flox/flox^ MEFs 4 days after infection with AdBlank or AdCre from 3 independent experiments. White lines denote examples of linear fluorescence intensity measurements as shown in Fig. 4f. Scale bar = 50 μm. (**f**) Fluorescence intensity profiles of CREB3 in *Lmna*^flox/flox^ MEFs infected with AdBlank or AdCre. The directionality of the fluorescence intensity measurements is from the cytoplasm (cyto) to the nucleus (nucl). n=number of nuclei measured. (**g**) Immunoblot of CREB3, LMNA, and GAPDH in *Lmna*^flox/flox^ MEFs 4 days after infection with AdBlank (bl) or AdCre (cre). Black arrowheads denote CREB3 and its cleavage. Gray arrowheads denote lamins A and C. Asterisk denotes non-specific band. (**h**) (Top left) Representative immunoblot of CREB3 and GAPDH in primary adult CMs isolated from CM-CreTRAP: *Lmna*^flox/flox^ mice treated with vehicle (veh) or Tam. 2w and 4w denote weeks post Tam treatment. Numbers on top of blots denote individual heart samples. Black arrowheads denote CREB3 and its fragments. Remaining graphs show qPCR analyses probed for *Arf4*, *Trappc13*, *Tnfrsf1b*, and *Tnfrsf10b*. *, **, and *** denote p < 0.05, p < 0.005, and p < 0.0005, respectively, using one way ANOVA with Dunnett’s post hoc with mean value for veh used as the reference control. n=3-5. (**i**) Confocal immunofluorescence images of CREB3 and DAPI counterstain on primary adult CMs isolated from CM-CreTRAP: *Lmna*^flox/flox^ mice treated with vehicle or 2 weeks post Tam. Two examples of 2 week post Tam data are shown. Scale bar = 10μm. (**j**) Quantification of CREB3 localization derived from the immunofluorescence data shown in Fig. 4i. n denotes number of nuclei analyzed. All error bars in this figure = SEM.

The swollen golgi stacks at the perinuclear space are reminiscent of a similar stress response induced by a ionophoric antibiotic monensin, which acts by collapsing Na^+^ and H^+^ gradients, causing golgi dilation(*51*). As previously indicated, prior studies have shown that monensin-induced golgi stress triggers the ATF4-CHOP axis of the UPR without the other two (XBP-1 and ATF6)(*49, 50*). Similar molecular expression profiles were observed in the hearts of our CM-*Lmna* deleted mice, strongly indicating that golgi stress response underlies the selective ATF4-CHOP activation. To confirm this and to establish causality, we utilized *in vitro* culture models to recapitulate the *in vivo* phenotype.

Golgi stress induced by monensin can activate CREB3, an ER-resident protein belonging to a family of transcription factors that includes ATF6(*52*). In a similar fashion to ATF6, full length CREB3 is translocated to the golgi in response to stress where it undergoes proteolytic cleavage, producing a transcriptionally active transcription factor that enters the nucleus and transactivates the expression of *ARF4*, *TRAPPC13*, and *TNFRSF10A*(*53*). Following 2 μM monensin treatment, both mouse embryonic fibroblasts (MEFs) isolated from *Lmna*^flox/flox^ mice (Fig. 4b, left panel) and C2C12 myoblasts (Supplementary Fig. 8d) displayed CREB3 cleavage (from uncleaved ∼46 kDa protein to ∼38 kDa fragments) by 24 hr. Assessment of mRNA upregulation of genes induced by CREB3 in MEFs revealed that although *Arf4* levels were significantly elevated by 4 hr after monensin treatment (Fig. 4b, right panel), *Trappc13* only showed a trend towards significance (Supplementary Fig. 8e). Moreover, we interestingly found that no murine homologue for *TNFRSF10A* exists. Therefore, we measured other members of the TNF family of receptors *Tnfrsf1b* and *Tnfrsf10b* and observed that they were also significantly elevated by 4 hr post monensin treatment (Fig. 4b, right panel), indicating that CREB3 may mediate the broad transactivation of TNF receptor family members. Furthermore, consistent with the prior reports(*49, 50*), monensin robustly activated the ATF4-CHOP axis, without similar activation of *Atf6*, *Hspa5*, and *Xbp1* (as well as its splicing) whereas brefeldin A displayed no such selectivity (Fig. 4c, d).

Having established the molecular phenotypes from golgi stress induced by monensin, we then assessed whether *Lmna* deletion, in and of itself, can induce CREB3 activation. To achieve this, we deleted *Lmna* in *Lmna*^flox/flox^ MEFs using adenovirus carrying cre recombinase (AdCre) and assessed CREB3 cleavage and nuclear entry. Relative to MEFs infected with a blank adenovirus (AdBlank), those infected with AdCre displayed increased CREB3 localization in the nucleus (Fig. 4e, f and Supplementary Fig. 9 for uncropped micrographs). Notably, the CREB3 localization appeared to be coincident with the emergence of misshapen nuclei after 4 days post AdCre transduction of MEFs cultured in regular tissue culture plastic. Also concomitant with this timepoint (4 days post adenoviral transduction), we consistently detected the presence of CREB3 cleavage product in MEFs infected with AdCre but not in those infected with the blank virus (Fig. 4g). These results indicate that *Lmna* deletion alone can trigger CREB3 activation. To confirm that golgi-stress induced CREB3 is activated in the primary tissue from mice, we isolated adult CMs from Tam-treated CM-CreTRAP:*Lmna*^flox/flox^ mice and assessed for CREB3 activation. Similar to our immunoblot data (Fig. 3b), we observed lamin A/C depletion in adult CMs 2 weeks post Tam relative to vehicle-treated controls (Supplementary Fig. 10a). Lamin A/C loss appears to coincide with the incipient phase of golgi stress signaling as ∼50% of 2 week adult CMs displayed CREB3 cleavage whereas all 4 week samples showed CREB3 cleavage as well as significant upregulation of *Arf4*, *Tnfrsf1b,* and *Tnfrsf10b* but not *Trappc13* (Fig. 4h). We further probed for CREB3 localization by immunofluorescence (Fig. 4i). In adult CMs from vehicle-treated mice, CREB3 was predominantly perinuclear (91.1% of adult CM nuclei analyzed), which is consistent with ER localization (Fig. 4i, j, and see Supplementary Fig. 10b, c for uncropped and additional images, respectively). However, following *Lmna* deletion, we observed 3 distinct patterns; CREB3 remained perinuclear (21%), was intranuclear (26.9%), or not detected (52.1%), indicating that CREB3 was activated in a subset of CMs (Fig. 4i, j). The significance of the differential localization pattern is currently unclear. Although other golgi stress responses have been identified, such as those involving Golph3 (*54*), we did not observe its activation in monensin-treated MEFs as well as in the *Lmna*-deleted hearts (data not shown), indicating that either it’s not involved or not as prevalent as CREB3-mediated responses. Our results show that *Lmna* deletion in CMs leads to CREB3-mediated golgi stress and the activation of the ATF4-CHOP signaling axis.

### Regulation of UPR by MED25

Given that MED25 was shown to facilitate ATF6-mediated transcription of *Hspa5* in response to ER stress inducer thapsigargin(*47*), we hypothesized that MED25 may contribute to the observed selective activation of the ATF-CHOP axis. To test this hypothesis, as well as to determine the functional relevance of MED25 expression in this context, we employed muscle myoblast C2C12 culture model with short hairpin RNA (shRNA)-mediated silencing of *Med25*. Following MED25 depletion (Supplementary Fig. 11a), C2C12 cells subjected to thapsigargin-induced ER stress were assessed for transcriptional responses in UPR. Despite comparable upregulation of *Atf4* and *Ddit3* mRNA between both MED25-depleted vs blank control cells in response to thapsigargin, *Atf6* and *Hspa5* (but not *Xbp1*) levels were super-induced with MED25 depletion at the 24 hr timepoint (Fig. 5a left panel and Supplementary Fig. 11b). Of note, opposite pattern of expression for *Hspa5* were observed between the 3 and 24 hr timepoints; MED25 knockdown reduced *Hspa5* expression at the earlier timepoint, suggesting a biphasic expression (Fig. 5a left panel). This effect appears to be selective as other stress triggers such as glucose starvation (Fig. 5a right panel), brefeldin A (Supplementary Fig. 11c), and monensin (Supplementary Fig. 11d) exhibited no such differences, indicating that the regulatory control exerted by MED25 on UPR appears to be more complex than anticipated. In a reciprocal experiment, transient overexpression of MED25 dampened *Atf6*, *Hspa5*, and *Xbp1* (only at 3hr) expression in response to thapsigargin but not under glucose starvation (Fig. 5b). These studies demonstrate that MED25 suppresses the ATF6 axis of UPR in response to thapsigargin and brefeldin A, but not to glucose starvation and monensin.

**Fig. 5.**
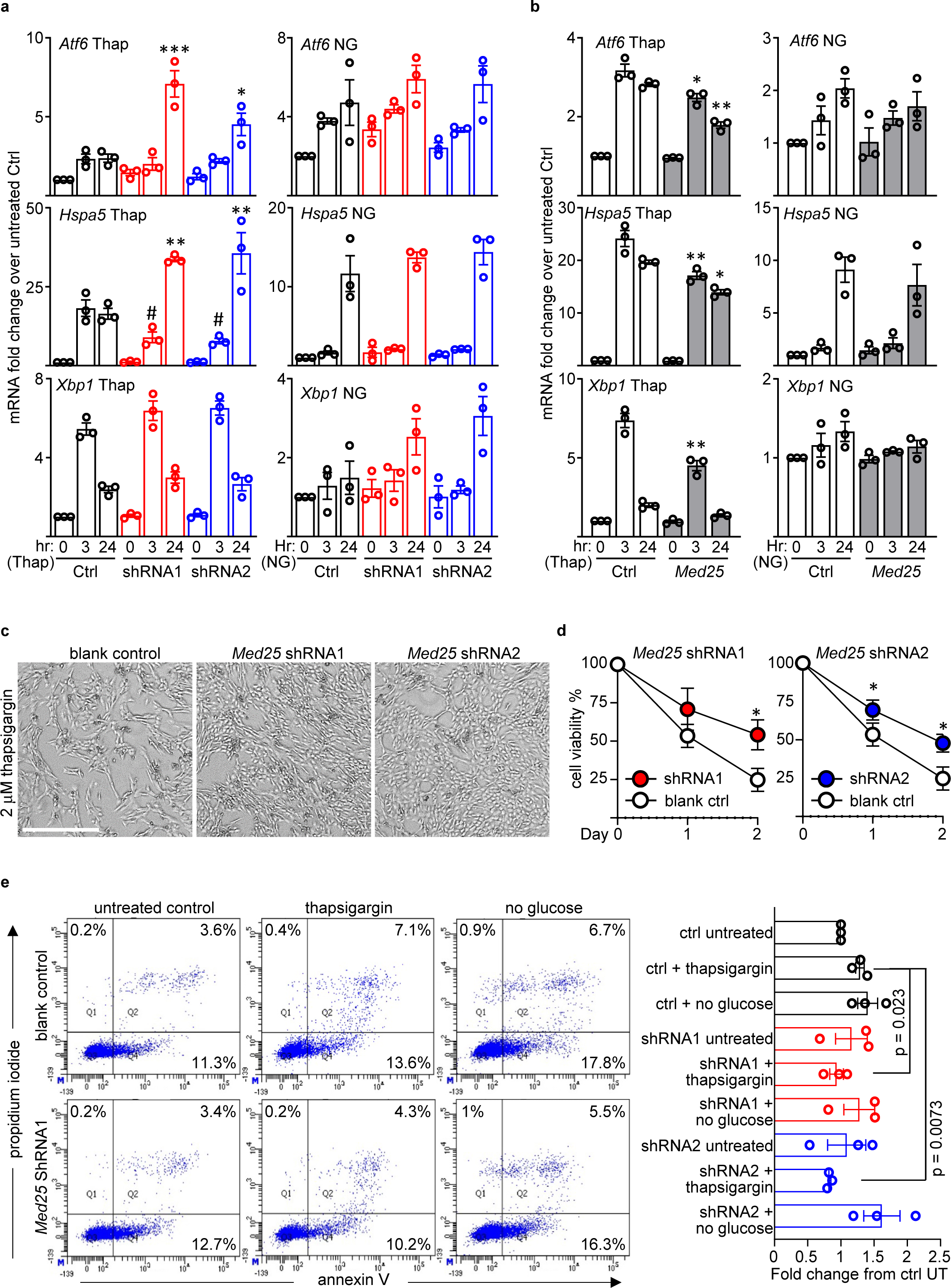
Selective regulation of UPR mediators by *Med25*. (**a**) qPCR analyses probing *Atf6*, *Hspa5*, and *Xbp1* mRNA in MED25-depleted C2C12 cells treated with 2 μM thapsigargin or starved of glucose for 3 and 24 hr. Fold change values were derived by setting untreated control as 1. NG denotes no glucose. *, **, and *** denote p < 0.05, p < 0.005, and p <0001, respectively, using one way ANOVA with Tukey correction with Ctrl 24 hr timepoint as reference. (**b**) qPCR analyses probing *Atf6*, *Hspa5*, and *Xbp1* mRNA in C2C12 cells overexpressing MED25 and treated with 2 μM thapsigargin or starved of glucose for 3 and 24 hr. Fold change values were derived by setting untreated control as 1. * and ** denote p < 0.005 and p < 0.0001, respectively, using one way ANOVA with Tukey correction relative to their respective Ctrl timepoints. (**c**) Representative phase contrast images of C2C12 cells with Med25 knockdown with two independent shRNAs and treated with 24 hr thapsigargin. Blank control denotes C2C12 infected with lentivirus generated using empty shRNA vector. (**d**) Viability of MED25-depleted C2C12 cells at 24 and 48 hr post 2 μM thapsigargin treatment. % viability was calculated relative to their respective cells treated with vehicle. * denote p < 0.05 using unpaired student’s T-test. Error bars = SD. (**e**) Representative images of AnnexinV/propidium iodide (PI) staining of MED25-depleted C2C12 cells treated with either 2 μM thapsigargin or starved of glucose for 24 hr. Bar graphs on right show fold change values, relative to “ctrl untreated”, of total cell death calculated as the sum of the 3 quadrants of AnnexinV/PI positive cells. P values were determine by one way ANOVA with Dunnett correction with ctrl + thapsigargin as control. Error bars = SEM. n=3 for all data in this figure.

The MED25 knockdown and the resulting super-induction of *Atf6* and *Hspa5*, which is considered to be cytoprotective, should theoretically enhance cell survival under thapsigargin treatment. Indeed, C2C12 cells with MED25 knockdown displayed enhanced cell viability (Fig. 5c, d). Annexin V/propidium iodide staining further confirmed that, although MED25 depletion itself seemed to slightly increase basal cell death, it conferred protection from thapsigargin-stimulated cytotoxicity but not from glucose starvation (Fig. 5e). These studies show that MED25 regulates cell survival and selective transcriptional responses to ER stress, particularly from those elicited by a calcium imbalance in the ER.

### Disruption of golgi underlies autophagic abnormalities

We and others previously showed that autophagy, a stress-responsive lysosomal recycling mechanism that maintains cellular homeostasis, is impaired in germline models of *LMNA* cardiomyopathy(*16, 17*). To determine whether a similar impairment in autophagy is present in our inducible CM-specific *Lmna* deletion model, we assessed the levels of lipidated LC3B (LC3B-II) produced during autophagosome formation. This data was analyzed in the context of the expression level of p62, an autophagosome-associated protein that is degraded when autophagosomes are fused with lysosomes in a process described as autophagic flux. The combination of these two protein levels can be interpreted to distinguish whether a reduction in LC3B-II is caused by impaired autophagosome formation or its rapid degradation, and conversely, an increase in LC3B-II as a rapid expansion of a productive autophagic response or a blockade in autophagic flux. In the hearts of CM-*Lmna* deleted mice, we observed a slight increase in both p62 and LC3B-II protein levels at 1 to 2 weeks post Tam treatment relative to vehicle controls (Fig. 6a, b). By 4 weeks, significant increases in p62 and LC3B-II were observed, which is indicative of autophagosome accumulation due to a blockade in autophagic flux.

**Fig. 6.**
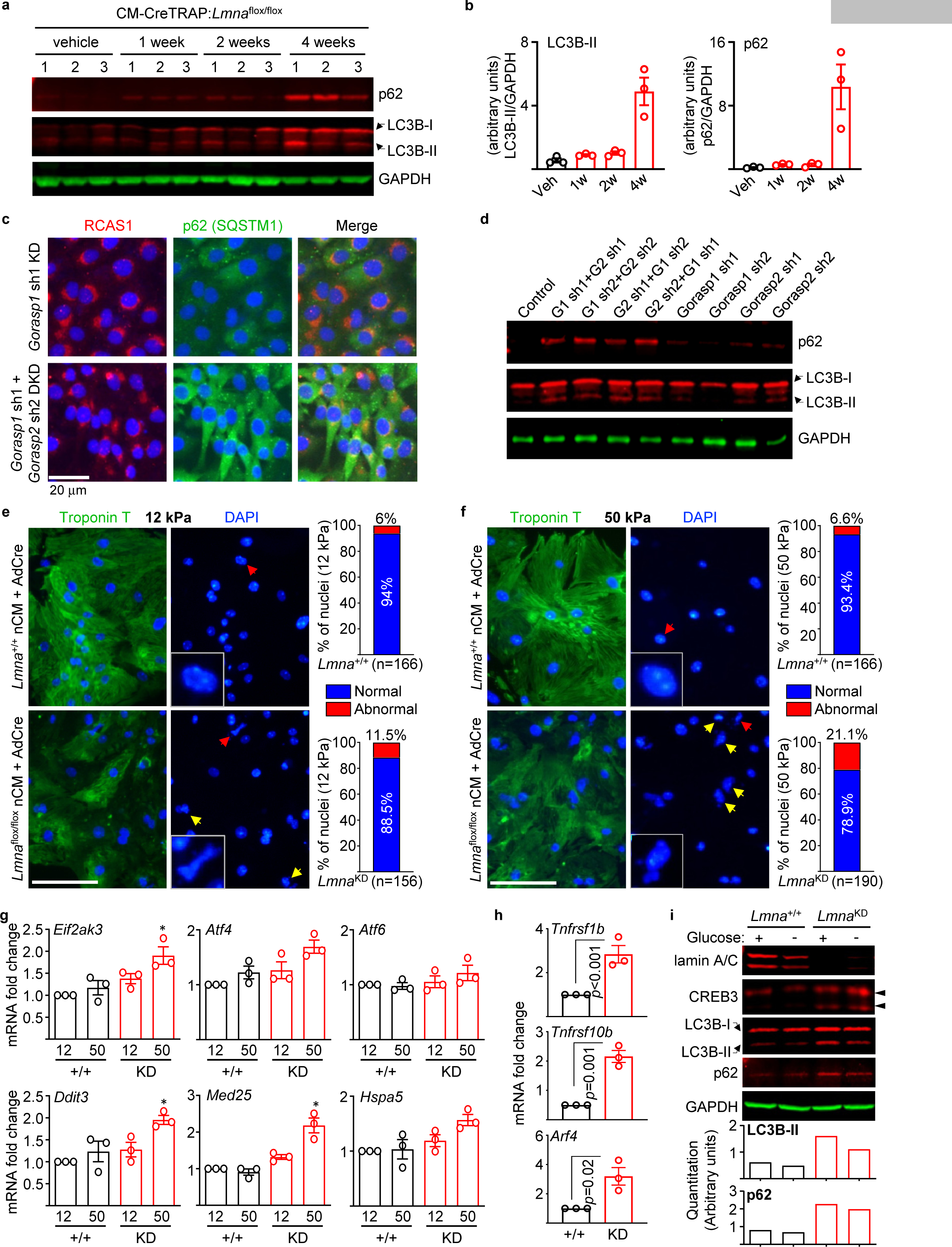
NE damage and golgi disruption underlie accumulation of autophagic markers. (**a**) Immunoblot of vehicle or Tam-treated CM-CreTRAP:*Lmna*^flox/flox^ heart extracts probed for p62 and lipidated LC3B [LC3B-II]. GAPDH was assessed as loading control. Weeks denote time since the last Tam dosing. Numbers on top of blots denote individual heart samples. (**b**) Quantitation of LC3B-II and p62 blots as shown in 6a normalized to GAPDH. (**c**) Representative immunofluorescence images of C2C12 cells with *Gorasp1* KD alone or *Gorasp1* and *Gorasp2* double KD (DKD) stained for RCAS1 and p62. Scale bar = 20 μm. (d) Immunoblot of C2C12 cells with *Gorasp1* KD alone or *Gorasp1*/*Gorasp2* DKD probed for p62 and LC3B-II. GAPDH was assessed as loading control. Control denotes C2C12 cells infected with lentivirus generated from blank plasmid **(e,f**) Troponin T and DAPI staining on nCMs isolated from *Lmna^+^*^/+^ and *Lmna*^flox/flox^ mice (*Lmna* KD) infected with adCre and cultured on 12 kPa matrix (**e**) or on 50 kPa matrix (**f**) for 48 hr. Yellow arrows denote ruptured and herniated nuclei and the red denoting nucleus in inset. Bar graphs show quantitation presented as % of normal or abnormal nuclei from total number of cells counted. Scale bar = 100 μm. (**g**) qPCR analysis of *Eif2ak3* (encoding PERK), *Atf4*, *Atf6*, *Ddit3* (CHOP), *Med25*, and *Hspa5* (Grp78) transcripts in nCMs isolated from *Lmna^+^*^/+^ (+/+) and *Lmna*^flox/flox^ mice (KD) infected with adCre and cultured on 12 and 50 kPa matrix for 48 hr. Fold change values were determined relative to *Lmna^+^*^/+^ nCM on 12 kPa as control set to 1. n = 3. * denotes p < 0.05 determine by one way ANOVA with Dunnett correction with *Lmna^+^*^/+^ nCM on 12 kPa as control. Error bars = SEM. (**h**) qPCR analysis of *Tnfrsf1b*, *Tnfrsf10b*, and *Arf4* transcripts on nCMs in nCMs isolated from *Lmna^+^*^/+^ (+/+) and *Lmna*^flox/flox^ mice (KD) infected with adCre and cultured on 50 kPa matrix for 48 hr. P values were derived using unpaired, two-tailed, Student’s t test. Error bars = SEM. (**i**) Immunoblot of lamin A/C, CREB3, LC3B, p62, and GAPDH on nCMs isolated from adCre-treated *Lmna*^+/+^ and *Lmna*^flox/flox^ mice (*Lmna*^KD^) cultured with and without glucose for 24 hr in 50 kPa matrix. Representative blot from n=3 experiments. Black arrowheads denote CREB3 and its fragments. Bottom panel shows quantitation of LC3B-II and p62 normalized to GAPDH.

Given that lysosomes originate from the golgi, we sought to determine whether direct disruption of the golgi apparatus will engender similar abnormalities in autophagy. To achieve this, we employed shRNA-mediated knockdown of golgi proteins GORASP1 and GORASP2 in C2C12 cells, as the depletion of both of these proteins were demonstrated to cause golgi disruption and disassembly(*55*). We generated double-knockdowns by a sequential delivery of shRNAs targeting *Gorasp1* and *Gorasp2* (Supplementary Fig. 12a) and then assessed for golgi by staining for RCAS1, a protein highly enriched in golgi(*56*). In C2C12 cells with a single knockdown (either *Gorasp1* or *Gorasp2*) as well as in blank controls (C2C12 infected with lentivirus carrying a blank vector), RCAS1 staining exhibited crescent perinuclear staining indicative of the golgi apparatus. In contrast, C2C12 with *Gorasp1* and *Gorasp2* double knockdown revealed a much smaller golgi footprint with many cells displaying reduced signals (Fig. 6c; uncropped images are shown in Supplementary Fig. 12b). These results are indicative of golgi disassembly and are consistent with the previously reported observations(*55*). We then assessed for p62 and LC3B-II and found that C2C12 cells with golgi disruption displayed increased accumulation of both of these proteins relative to singly targeted cells even under basal conditions (Fig. 6d). This accumulation is post-transcriptionally mediated as no change in the mRNA levels of *Sqstm1* (p62) and *Map1lc3b* (LC3B) were detected (Supplementary Fig. 12c). We also probed for transcriptional activation of genes encoding ATF4-CHOP and observed that golgi disruption did not lead to their increase (Supplementary Fig. 12c). These results demonstrate that p62 and LC3B-II accumulation results from golgi disruption that is, in and of itself, insufficient to trigger the ATF4- CHOP axis and presumably require additional stressor such as nuclear envelope damage.

### Physiological conditions recapitulate molecular pathogenesis from Lmna deletion

Physical tension felt by the nucleus is positively correlated with matrix stiffness(*57*). Given our hypothesis that nuclear rupture resulting from lamin A/C depletion causing collateral damage to the golgi in the perinuclear space, it is important to establish that the physical and molecular perturbations can be recapitulated under physiologically conditions in relevant cell types. Although we noted enhanced nuclear damage (and coincident CREB3 localization) in lamin A/C-depleted MEFs cultured on hard plastic, these conditions are not physiologically relevant. Therefore, we depleted lamin A/C in nCMs isolated from *Lmna*^flox/flox^ mice and cultured in either 50 kPa matrix, which is similar in stiffness to a fibrotic cardiac tissue, or 12 kPa matrix (normal adult myocardium)(*58, 59*). Following infection with AdCre and culture in a 12 kPa matrix, ∼11% of *Lmna*^flox/flox^ nCMs (denoted as *Lmna*^KD^) exhibited perturbed nuclei whereas only ∼6% of *Lmna*^+/+^ nCMs displayed similar abnormalities (Fig. 6e). However, a significantly higher percentage of *Lmna*^flox/flox^ nCMs displayed nuclear damage (∼21.1%) when cultured in a 50 kPa matrix compared to *Lmna*^+/+^ nCMs, which remained relatively steady at ∼6.6% (Fig. 6f). These features were associated with significant increases in transcripts encoding PERK and CHOP as well as *Med25* relative to *Lmna*^+/+^ nCMs (Fig. 6g). Assessment of golgi stress-associated transcripts *Tnfrsf1b*, *Tnfrsf10b*, and *Arf4* were significantly elevated in *Lmna*^KD^ nCMs relative to *Lmna*^+/+^ nCMs when cultured in 50 kPa (Fig. 6h). Moreover, *Lmna*^KD^ nCMs displayed increased accumulation of both LC3B-II and p62 levels irrespective of glucose starvation (Fig. 6i). Similarly, CREB3 cleavage was observed in *Lmna*^KD^ nCMs irrespective of glucose presence but not in *Lmna*^+/+^ nCMs (Fig. 6i). Our results indicate lamin A/C-depletion in nCMs under physiologically relevant conditions recapitulate the observed phenotypes *in vivo*.

### Enhancing autophagy or modulating ER stress preserves cardiac function

We and others previous demonstrated that reactivation of autophagy by rapamycin and its analog temsirolimus mitigates the disease in germline models of *LMNA* cardiomyopathy(*16, 17*), indicating a protective role of autophagy in this setting. Our current study suggests damage in the perinuclear space emanating from nuclear fragility underlies the initial stressor that elicits a protective autophagy response and when impaired, the damage to the CMs is exacerbated (*60*). Given the similar impairment in autophagy observed in our CM-specific model, we sought to perform pharmacological intervention studies aiming to mitigate stress signaling emanating from CM-specific lamin A/C depletion. Following CM-*Lmna* deletion in the adult heart, we administered either metformin (to stimulate autophagy) or azoramide (ER stress modulator)(*61*), as specific golgi stress modulators have yet to be identified. Azoramide is a small molecule demonstrated to improve ER protein-folding ability by stimulating multiple chaperone expression in a dose-dependent fashion(*61*). Because high dose azoramide administration (150 mg/kg) can enhance antidiabetic effects and glucose homeostasis(*61*), we decided on a far lower dose of 50 mg/kg (Fig. 7a). Given that the therapy was initiated prior to detectable cardiac function decline, our strategy represents a preventative therapy. Although no significant differences in cardiac function were observed at 2 weeks post Tam with or without metformin/azoramide relative to vehicle controls, these drugs significantly reduced both the systolic and the diastolic ventricular dimensions by 4 weeks, which resulted in preserved ejection fraction (Fig. 7b). Histopathology revealed that while the extent of cardiac fibrosis was similar between Tam versus Tam with metformin, the myocardial tissue disarray, cytoplasmic lysis and vacuolation were less obvious (Fig. 7c and Supplementary Fig. 7a). Conversely, despite the similar preservation of cardiac function, hearts from azoramide-treated mice were phenotypically different.

**Figure 7.**
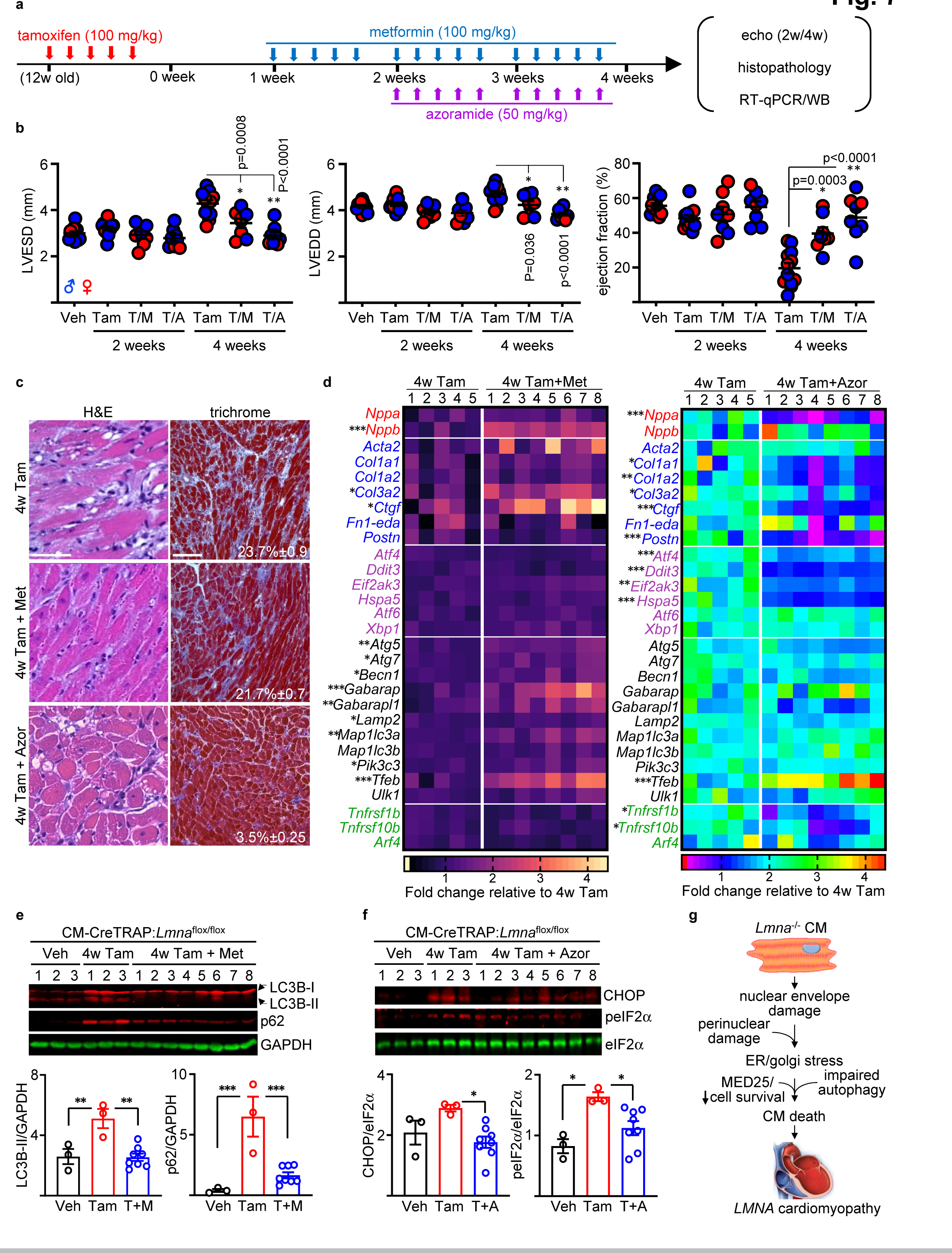
Enhancing autophagy or modulating ER stress attenuates cardiomyopathy development. (**a**) Schema of Tam, metformin, and azoramide dosing schedule. “12w old” denotes 12-week-old mice. (**b**) Echo on CM-CreTRAP:*Lmna*^flox/flox^ mice at 2 and 4 weeks after Tam alone (Tam), Tam + metformin therapy (T/M), or Tam + azoramide (T/A). LVESD and LVEDD denote left ventricular end systolic and diastolic diameters, respectively. p values were obtained using one-way ANOVA with Tukey correction. Error bars = SEM. (**c**) H&E and Masson’s Trichrome staining of the heart sections from CM- CreTRAP: *Lmna*^flox/flox^ mice at 4 weeks post tamoxifen (4w Tam), with metformin (4w Tam + Met) or with azoramide (4w Tam + Azor). Scale bar = 100 μm. (**d**) Heat map of expression analyses on natriuretic peptide precursors (red), profibrotic (blue), ER stress- (purple), autophagy (black), and golgi stress (green)-related mRNA expression in the hearts from CM-CreTRAP:*Lmna*^flox/flox^ mice at 4 weeks post Tam treatment alone (4w Tam), with metformin (left panel, 4w Tam+Met), or with azoramide therapy (right panel, 4w Tam+Azor). Numbers on top of heatmap denote individual heart samples. Color gradient scales and values on the bottom denote fold change over mean value of 4w Tam from the 5 biological replicates. p values were obtained using unpaired, two-tailed Student’s t test. * = p < 0.05, ** = p < 0.01, *** = p < 0.001. (**e**) Immunoblot on heart extracts from CM-CreTRAP:*Lmna*^flox/flox^ mice at 4 weeks post vehicle, Tam treatment alone, and with metformin therapy and probed for p62, LC3B, and GAPDH. Numbers on top of blots denote individual heart samples. Bottom panel shows quantitation of blots normalized to loading controls. p values were obtained using one way ANOVA with Tukey correction. * = p < 0.05, ** = p < 0.01, *** = p < 0.001. Error bars = SEM. (**f**) Immunoblot similar to above but with azoramide therapy and probed for CHOP, phospho-eIF2α, and total eIF2α. (**g**) Working hypothesis for *LMNA* cardiomyopathy pathogenesis. See text for details.

Although vacuolation was still obvious, we observed significantly reduced fibrosis (Fig. 7c and Supplementary Fig. 7a). These histological phenotypes were further supported by mRNA expression profiling analyses. We interrogated the expression of genes involved in several key cellular responses; cardiac stress adaptation- (red), profibrotic- (blue), UPR- (purple), autophagy- (black), and golgi stress- (green) related (Fig. 7d). Although several genes were enhanced by metformin relative to Tam alone, by far the largest number of significantly enhanced genes was in the autophagy response (left panel on Fig. 7d and Supplementary Fig. 7c). Alternatively, azoramide treatment predominantly reduced profibrotic genes as well as the UPR genes tested relative to Tam alone, although *Atf6* and *Xbp1* were not significantly altered (right panel on Fig. 7d and Supplementary Fig. 7c). Notably, *Tnfrsf1b* and *Tnfrsf10b* expressions were significantly reduced with the azoramide treatment but not with metformin despite the similar improvement in heart function, suggesting that azoramide may provide partial modulation of golgi stress (Fig. 7d). At the protein level, metformin treatment reduced both LC3B-II and p62 levels, indicative of restored autophagy (Fig, 7e). Analogously, azoramide treatment reduced phospho-eIF2α (upstream activator of PERK) as well as CHOP expression (Fig. 7f), confirming the effects of azoramide treatment at the protein level. Altogether, these results indicate that CM-specific *Lmna* deletion leads to impaired autophagy in the heart and either restoring autophagy or alleviating its need by dampening ER stress can delay *LMNA* cardiomyopathy.

## Discussion

It is well appreciated that lamin A/C maintain structural support of the nucleus, but how nuclear deformities resulting from *Lmna* mutations underlie pathology in a complex organ such as the heart is less well understood. In the current study, we found that adult CM-specific lamin A/C depletion caused rapid development of cardiomyopathy with accompanying pathological fibrosis (Fig. 7g). Prior to cardiac function decline, NE damage elicited CREB3-mediated golgi stress response and selective activation of the ATF4-CHOP axis. We further link Med25, a member of the mediator complex, to selective UPR activation in the *LMNA* cardiomyopathy pathogenesis, which sensitized cells to ER stress. Lastly, we show that golgi disruption caused p62 and LC3B-II accumulation and the administration of autophagy inducing drug metformin enhanced their clearance and ameliorated the disease. Likewise, a similar benefit to the cardiac function was achieved by UPR modulator azoramide.

To our knowledge, this is the first study demonstrating a connection between golgi stress as a consequence of *Lmna* deletion in CMs. Furthermore, the accumulation of p62 and lipidated LC3B as a direct consequence of golgi disruption is a previously undescribed observation. We assert that the interplay of stress responses emanating from NE and perinuclear organelle damage are at the nexus of the pathogenesis of *LMNA* cardiomyopathy. Golgi was first hypothesized to be implicated in laminopathies based on the observation that MEFs from *Lmna^-/-^* mice displayed Sun1 accumulation in the golgi(*62*). Although it is plausible that Sun1 mislocalization contributes to the molecular pathogenesis, our data suggest that ionic/pH changes more likely underlie the dilation and the subsequent fragmentation of golgi. This is based on 1) the observation that golgi damage and the resulting stress signaling in the CMs of *Lmna-*deleted hearts are similar to those induced by monensin, which acts by collapsing the Na^+^ and H^+^ gradients(*51*) and 2) the existence of tightly regulated Na^+^ and H^+^ gradients across the nuclear envelope maintained by Na^+^-K^+^-ATPase and vacuolar H^+^ ATPase, respectively(*63*).

An additional novel component of our study is the identification of Med25 upregulation in *LMNA* cardiomyopathy. Although the exact mechanisms remain to be elucidated, Med25 selectively suppressed cytoprotective axes of UPR in response to thapsigargin but not to other ER stressors such as brefeldin A and glucose starvation as well as to golgi stressor monensin. Interestingly, golgi stress regulator CREB3 and its related family member CREB3L2 were specifically activated in response to monensin and brefeldin A but not thapsigargin(*64*). An obvious explanation may be that transcriptional regulators governing UPR and possibly golgi stress are context-specific such that Med25 activity is further modulated in the presence of secondary calcium signaling that occurs with thapsigargin. These observations highlight a complex molecular circuity governing the various UPR/golgi stress pathways. It also remains to be established whether ER stress (and therefore complete UPR activation) precedes golgi stress or whether they are mutually exclusive manifestations. It is attractive to hypothesize that they are connected in a sequential manner in which the NE damage eventually leading to golgi stress is a “commitment to cell death” signal, which is consistent with the selective activation of the ATF4-CHOP.

Some limitations are worth noting. Although metformin can stimulate autophagy(*65*), it also exhibits pleiotropic effects in an organ-specific fashion. For example, metformin blocks hepatic gluconeogenesis, activates AMPK in striated muscle, and alters the gut microbiota in the intestine(*66*). Therefore, while the metformin therapy resulted in transcriptome changes indicative of increased autophagy in our study, the correlating cardiac protection observed may have benefitted from the aforementioned systemic effects of metformin. Same can be said for azoramide as its use at higher concentrations can have metabolic benefits as mentioned previously. Therefore, although the positive effects of these drugs on the heart function are only suggestive of autophagy blockade and ER stress involvement in the pathogenesis of *LMNA* cardiomyopathy, they still establish translational potential that can be realized into therapeutic strategies for a disease that, to date, has no effective targeted therapy. It’s also worth noting the limitation of our 2-dimensional (2D) stiffness culture model to recapitulate the *in vivo* condition. In the 2D culture, the nuclear damage likely result from compressive forces generated by cell flattening(*19*). However, forces applied to CMs in an intact heart would likely be more complex than a unidirectional compressive force. Nevertheless, given the recapitulation of nuclear abnormalities in 2D culture, we posit that the main driver of the observed nuclear damage is due to the lack of lamin A/C expression despite the noted differences in the forces felt by the CMs.

## Materials and Methods

### Animals

All animal procedures were approved by the Institutional Animal Care and Use Committee of Thomas Jefferson University. All methods adhered to the NIH Guide for the Care and Use of Laboratory. *Lmna*^flox/flox^ mice(*34*), obtained from The Jackson Laboratory on a mixed background, were backcrossed to C57BL/6J for at least 8 generations and genotyped as instructed by the distributor. CM-CreTRAP transgenic mice were generated in a C57BL/6 background by Cyagen Biosciences. Sequences encoding EGFP and murine RPL10a was obtained and fused according to Heiman et al(*23*) using In-Fusion HD (Takara Bio cat# 011614). The EGFP-L10a fusion construct was incorporated downstream of CreERT2(*67*) (Addgene #14797) linked by an IRES (Addgene #12254) while simultaneously being subcloned into pPSKII-MHC-GH(*68*) downstream of the *Myh6* (α- myosin heavy chain) promoter and upstream of a human growth hormone polyadenylation signal using In-Fusion HD (Takara Bio cat# 011614). The final transgenic targeting construct was verified by DNA sequencing. Genotyping was performed by standard PCR using genomic DNA isolated from tail clippings and primers indicated in the Supplementary Table 3. Mice were fed a chow diet and housed in a disease-free barrier facility with 12/12 hr light/dark cycles. Tamoxifen (MilliporeSigma cat# T5648) was dissolved in corn-oil (MilliporeSigma cat# C8267) and injected intraperitoneally. Intraperitoneal delivery of Metformin HCl at 100 mg/kg (MilliporeSigma cat# PHR1084) was freshly prepared by dissolving in H_2_O and filter-sterilized prior to injection. Azoramide (Cayman Chemical cat# 18045) was dissolved in DMSO, filtered-sterilized, and injected at 50 mg/kg.

### Human samples

Human heart tissue from patients with *LMNA* cardiomyopathy was obtained at the time of heart transplantation and myocardium from age- and sex-matched nonfailing hearts was obtained from brain-dead organ donors. Use of human heart tissue for research was approved by the University of Pennsylvania Institutional Review Board and the use of hearts from brain-dead organ donors for was approved by the Gift-of-Life donor Program in Philadelphia, PA. Written informed consent transplant for research use of heart tissues was obtained prospectively from transplant recipients and from next-of-kin in the case of organ donors. In all cases, hearts were arrested in situ with ice-cold, high-potassium cardioplegia, excised from the body, and transported to the lab in ice-cold Krebs-Henseleit Buffer. Transmural samples taken from the left ventricular free wall were flash frozen in liquid nitrogen and stored at -80oC until tissue analyses were performed.

### Primary CM isolation and Lmna deletion

Murine nCMs were isolated from the ventricles of 1-2 days old wildtype C57BL/6 and *Lmna*^flox/flox^ mouse pups using MACS neonatal heart dissociation kit according to the manufacturer (Miltenyi Biotec cat# 130-098-373). Following digestion, the single cell suspension was pre-plated for 30 minutes to selectively remove non-myocyte populations. The non-attached phase was recollected and replated in plates coated with 10 μg/ml laminin (Life Technologies cat# 23017015) in DMEM supplemented with 10% fetal bovine serum and antibiotic/antimycotic solution. For experiments with culture in varying stiffness, nCMs cells were plated onto 10 μg/ml laminin-coated wells with 12 kPa or 50 kPa hydrogels (Matrigen cat# SW12-EC-12 PK and SW12-EC-50 PK). To prevent the potential growth of non-myocyte cells, the culture media was also supplemented with 100 μM 5-bromo-2-deoxyuridine (BrdU) and 10 µM cytosine arabinoside (Ara-C). The cells were maintained at 37 °C in 95% humidity with 5% CO_2_ concentration. The cells were left to attach overnight after which the media (DMEM+10% FBS+ 10 µM Ara-C) was replenished after rinse with PBS to remove dead cells. Adult CM isolation was performed as described here(*69*) using constant pressure Langendorf perfusion system using Liberase TH (MilliporeSigma cat# LIBTH-RO) at 25 μg/ml. To delete *Lmna in vitro*, adenovirus carrying mCherry (AdBlank) or mCherry-Cre (AdCre) (Vector Biolabs cat# 1767 and 1773, respectively) were used at 50 MOI. Similar infection efficiencies were confirmed by mCherry.

*C2C12 culture, shRNA knockdown, stress induction, and cell death analysis.* C2C12 cells (American Type Culture Collection) were maintained in DMEM supplemented with 10% FBS (VWR cat# 89510-186) at 37°C with 5% CO_2_ and subcultured at ∼60-70% confluency. Viral packaging cell line 293T cells were maintained in the same media. For stable knockdown of *Med25*, we used two independent shRNAs in pLKO.1 lentiviral vector identified from murine *Med25* shRNA (MilliporeSigma cat# SHCLNG-NM_029365) with the sequences GCAGCTGTTCGATGACTTTAA (shRNA1) and TGCAGCTGTTCGATGACTTTA (shRNA2). For stable knockdown of *Gorasp1* and *Gorasp2*, we used two independent shRNAs for each in pLKO.1 lentiviral vector with the following sequences: CTCTGAAGCTGATGGTGTATA (shRNA1) and ACCTCACAACTTACTGCCTTT (shRNA2) for *Gorasp1* and GCTATGGTTATTTGCACCGAA (shRNA1) and CCCTGTCATGACTACTGCAAA (shRNA2) for *Gorasp2*. The lentiviral vectors were co-transfected into 293T cells with the packaging vectors pCMV-dR8.2 dvpr and pCMV-VSV-G (Addgene). Virus-infected cells were selected with 2 μg/ml puromycin. To induce stress, the cells were treated with 2 μM thapsigargin (Cayman Chemical cat# 10522), 20 μM brefeldin A (Cayman Chemical cat# 11861), 2 μM monensin (Cayman Chemical cat# 16488), or starved of glucose as previously described(*70*). Cell viability was determined by trypan blue exclusion assay. Annexin V/propidium iodide staining was performed using the Dead Cell Apoptosis Kit (ThermoFisher cat# V13242) according to the manufacturer’s instructions, sorted on BD Celesta and analyzed using FlowJo software.

### RNA isolation and RT-qPCR

Total RNA from tissues were isolated using Direct-zol RNA kit (Zymo Research cat# R2053) with a minor modification. Tissues were homogenized in TRIzol (Zymo Research cat# R2050-1-200) and the aqueous phase containing the RNA fraction was separated by adding chloroform (20% volume of TRIzol). The aqueous fraction (being careful to avoid genomic DNA interface) was mixed with 100% molecular grade ethanol at 1:1 ratio and then further processed using the Direct-zol RNA kit according to the manufacturer’s instructions. cDNA were generated from RNA (100ng for translating RNA, 1µg for total RNA) primed with a 1:1 ratio of random hexameric primers and oligodT using RevertAid RT kit (ThermoFisher Scientific cat# K1691). qPCR was performed on an QuantStudio5 qPCR system (Life Technologies) using PowerUP SYBR-green (ThermoFisher Scientific cat# A25743). *Gapdh* was assessed to ensure fidelity of enzymatic reactions and used as internal controls to normalize qPCR results. Fold-changes in gene expression were determined by the ΔΔCt method(*71*) and presented as fold-change over negative controls. For *Xbp1* splicing, cDNA generated were amplified using standard thermocycler GeneAmp PCR system 9700 (Applied Biosystems) and the amplicons were run on 2% agarose gel. A complete list of primer sets used in the study are provided in the Supplementary Table 3.

### Protein extraction, immunoblot analysis

Heart tissues were homogenized in ice-cold radioimmunoprecipitation assay (RIPA) buffer (MilliporeSigma cat# R0278) with Pierce Protease Inhibitor cocktail (ThermoFisher Scientific cat# A32963) and 1 mM sodium vanadate (MilliporeSigma cat# S6508). Following brief sonication (Dismembrator Model F60, ThermoFisher Scientific), the samples were prepped in Laemmli buffer after which 15 to 30 µg of protein extract was loaded for SDS-PAGE. Antibodies and the dilutions used in the study are provided in the Supplementary Table 3. Proper loading was initially confirmed by Ponceau S then by probing with GAPDH antibodies or otherwise indicated. Image capture was performed by Odyssey® Fc Imaging System and blot quantification was performed using Image Studio software (LI-COR Biosciences).

### Translating ribosome affinity purification and RNAseq

For TRAP, we processed the ventricular tissue exactly as described previously(*23*). 12- week-old mice were treated with either vehicle or Tam as described in Fig. 4a. 2 weeks after Tam (or vehicle) dosing, ventricular tissue were harvested and translating mRNAs purified from n = 3 biologically independent samples. Sequencing libraries were prepared from RNA samples using QuantSeq (Lexogen) and sequenced in a 75bp single end run on NextSeq 500 sequencer. RNAseq data was aligned using STAR algorithm against mm10 mouse genome version and RSEM v1.2.12 software was used to estimate read counts and FPKM values using gene information from Ensembl transcriptome version GRCm38.89. Raw counts were used to estimate significance of differential expression difference between any two experimental groups using DESeq2 (ref3). Overall gene expression changes were considered significant if passed FDR<5% threshold. Sequencing and data analysis were performed by Wistar Genomics and Bioinformatics Facilities. The data was submitted to NCBI GEO database under the accession number GSE185620.

### Transmission electron microscopy

Heparinized animals were euthanized and the excised hearts were cannulated through aorta on Langendorff apparatus. The hearts were initially perfused with Tyrode solution for 3 min (in mmol/l: 135 NaCl, 5.4 KCl, 5 MgCl_2_, 1 CaCl_2_, 0.33 NaH2PO_4_, 10 HEPES, pH 7.3), followed by perfusion with calcium free Tyrode solution (same as above except for 0.02 CaCl_2_), and finally by perfusion fixation with 2.5% glutaraldehyde) in 0.15 M sodium cacodylate buffer (pH 7.4). Small pieces from left ventricle were cut (∼1-3 mm^3^), postfixed overnight in 4°C in 2% osmium tetroxide partially reduced by 0.8% K4Fe(CN)6 in 0.15 M Na-cacodylate buffer. Samples were contrasted en bloc with 1% uranylacetate in diH2O, dehydrated in graded series of acetone, and embedded in Spurr’s resin. Longitudinal, ultrathin sections (65–80 nm) were cut from the resin-embedded blocks with a diamond knife (Diatome-US, USA) using a Leica UCT ultramicrotome and caught on copper grid covered with formvar film. Images of longitudinal oriented CMs were obtained via an FEI Tecnai 12 TEM fitted with an AMT XR-111 10.5 Mpx CCD camera at 3,200 - 15,000x magnification (80 kV). For quantitation of damaged nuclei (as determined by ruffled NE or those showing NE breakdown coupled with 100 nm vesicles present in the proximity) and golgi dilation (determined as doubling/tripling of the golgi stack width coupled with fragmentation), they were counted by a blinded observer unaware of the treatment conditions. For abnormal nuclei, the data is presented as a % of abnormal/normal nuclei from vehicle-treated (n=45), 2 weeks post Tam (n=31), and 4 weeks post Tam-treated mice (n=27) with n denoting the total number of unique nuclei identified from EM images from 2 independent hearts per group. For golgi dilation, the data is presented as a % of unique nuclei with dilated golgi in the perinuclear space.

### Light microscopy and histopathological analysis

H&E, Masson’s Trichrome, and Picro-Sirius Red staining were performed by Translational Research & Pathology Shared Resources (Thomas Jefferson University, PA) using standard methods. For WGA staining, frozen heart sections (8 µm) were fixed in ice-cold 4% paraformaldehyde (PFA) for 15 minutes, washed with 0.25% PBS-tween, permeabilized with 0.2% Triton X-100 for 5 minutes, and blocked with 1% bovine serum albumin in PBS-T for 1 hr. The sections were stained with Alexa Fluor™ 488 conjugated WGA solution (Invitrogen cat# W11261) at 5 μg/ml in HBSS for 15min at room temperature. Nuclei were counterstained with DAPI and the sections were mounted on coverslips with Prolong^TM^ Diamond Antifade Mountant (Invitrogen cat# P36961). For immunofluorescence, cells/tissue sections were fixed in ice-cold methanol:acetone (3:1) and processed using standard methods with antibodies at listed concentrations in Supplemental Table 4. All image analysis was performed using ImageJ 2.0 software(*72*). For myocyte cross-sectional area by WGA, the CMs were clearly distinguished from one another by their distinct cell membranes stained with WGA. Partial or incomplete cells were excluded from the analysis. 3 images were taken per section and area of 30 individual CMs were measured in total. Values were expressed as the average of area/myocyte in microns. For quantification of fibrosis by Masson’s Trichrome staining, percent fibrotic area was calculated by dividing blue pixels in each image by total number of pixels taken up by the heart section in indicated number of independent images per condition. Quantification of % nCMs containing perturbed nuclear shape was performed by dividing the number of cells containing abnormal nuclei by the total number of cells (∼150 to 190 cells per condition) across 6 images per condition from two independent experiments. Fluorescence intensity profiles of CREB3 traversing the NE were determined using the ImageJ(*72*) as previously described(*73*). Linear per-pixel fluorescence intensity of CREB3 was measured from the indicated number of cells from three separate experiments.

### Cardiac function

Mice were anesthetized with 1-2% isoflurane and placed on a stereotactic heated scanning base (37°C) attached to an electrocardiographic monitor. Left ventricular function was determined using a Vevo 2100 imaging system (VisualSonics) equipped with the MS550D transducer (22–55 MHz). Parameters were measured for at least three cardiac cycles between the heart rate of 400 to 500 beats per minute. A minimum of three separate M-mode images were used to derive the cardiac cycle parameters. An echocardiographer, blind to mouse genotype and/or treatment groups, performed the examinations and interpreted the results using AutoLV Analysis Software (VisualSonics).

### Statistical analysis

Graphpad (Prism 9) was used to perform statistical analyses. Statistical significance of binary variables was determined by a 2-tailed, unpaired Student’s *t-*test with a value of *p* < 0.05 considered significant. Statistical significance of three or more comparisons was determined by one way ANOVA with post-hoc Tukey error correction for multiple comparisons in which mean values were compared to all other means. For comparisons in which three or more means were compared to a defined control, one way ANOVA with Dunnett’s multiple comparison test was used. Values with error bars shown in figures are means ± SEM unless indicated otherwise. Sample sizes are indicated in the figure legends. *P* < 0.05 was considered significant.

## Acknowledgments

We thank Drs. Raymond Penn, Shey-Shing Sheu, and Tonio Pera for insightful discussions.

## Author contributions

K.S., E.P., and J.C.C. generated the hypotheses and designed experiments. K.S., E.P., D.M., J.S, and J.C.C. performed experiments, generated data in all figures and Supplementary Data. N. W. and E.P performed and analyzed echocardiography experiments. Z. Z. Performed and analyzed pathohistological analyses. K. B. M. provided and helped analyze human patient samples. A.K. performed and analyzed TRAP sequencing experiments. T.S., Z.N., and G.C. performed and analyzed TEM experiments. K.S., E.P., and J.C.C. wrote the manuscript. All authors carefully reviewed and edited the manuscript prior to submission.

## Data and materials availability

All data associated with this study are available in the main text or the Supplementary Materials. Heart tissue from human subjects with *LMNA* cardiomyopathy as well as the sex- and age-matched control samples were obtained through Uniform Biological Materials Transfer Agreement with The Trustees of the University of Pennsylvania. The samples were collected de-identified and not specifically for the proposed research by interacting with living individuals. TRAP GFP antibodies were obtained through MTA with Bi-Institutional Antibody and Bioresource Core Facility. TRAP sequencing data is available in NCBI GEO database under accession number GSE185620.

## Supplementary Materials

Supplementary Fig. 1. Generation and EGFP-L10a expression in CM-CreTRAP mice.

Supplementary Fig. 2. CM-specific depletion of lamin A/C.

Supplementary Fig. 3. CM-specific depletion of lamin A/C causes cardiac remodeling.

Supplementary Fig. 4. Cardiac perturbations in CM-specific *Lmna-*deleted mice.

Supplementary Fig. 5. Quality control data for TRAP sequencing.

Supplementary Fig. 6. Validation of MED25 and MED6 expression in *Lmna*-deleted hearts.

Supplementary Fig. 7. Perinuclear abnormalities in hearts of CM-specific *Lmna*-deleted mice.

Supplementary Fig. 8. Golgi abnormalities in hearts of CM-specific *Lmna-*deleted mice. Supplementary Fig. 9. Uncropped images shown in Fig. 4e.

Supplementary Fig. 10. CREB3 localization to the nucleus in adult CMs with *Lmna* deletion.

Supplementary Fig. 11. *Med25* function in C2C12 cells.

Supplementary Fig. 12. Golgi disruption in C2C12 culture model.

Supplementary Fig. 13. Characterization of Tam-treated CM-CreTRAP:*Lmna*^flox/flox^ mice treated with metformin or azoramide.

Supplementary Table 1. Echocardiography table for Fig. 1d.

Supplementary Table 2. Echocardiography table for Fig. 7b. Supplementary Table 3. Primers used in the manuscript.

Supplementary Table 4. Antibodies used in the manuscript.

