## Supplemental Figures for "Perinuclear damage from nuclear envelope deterioration elicits stress responses that contribute to *LMNA* cardiomyopathy"

Kunal Sikder *et al.*

**This PDF file includes:**

- Supplementary Fig. 1 - 13
- Supplementary Table 1 - 4

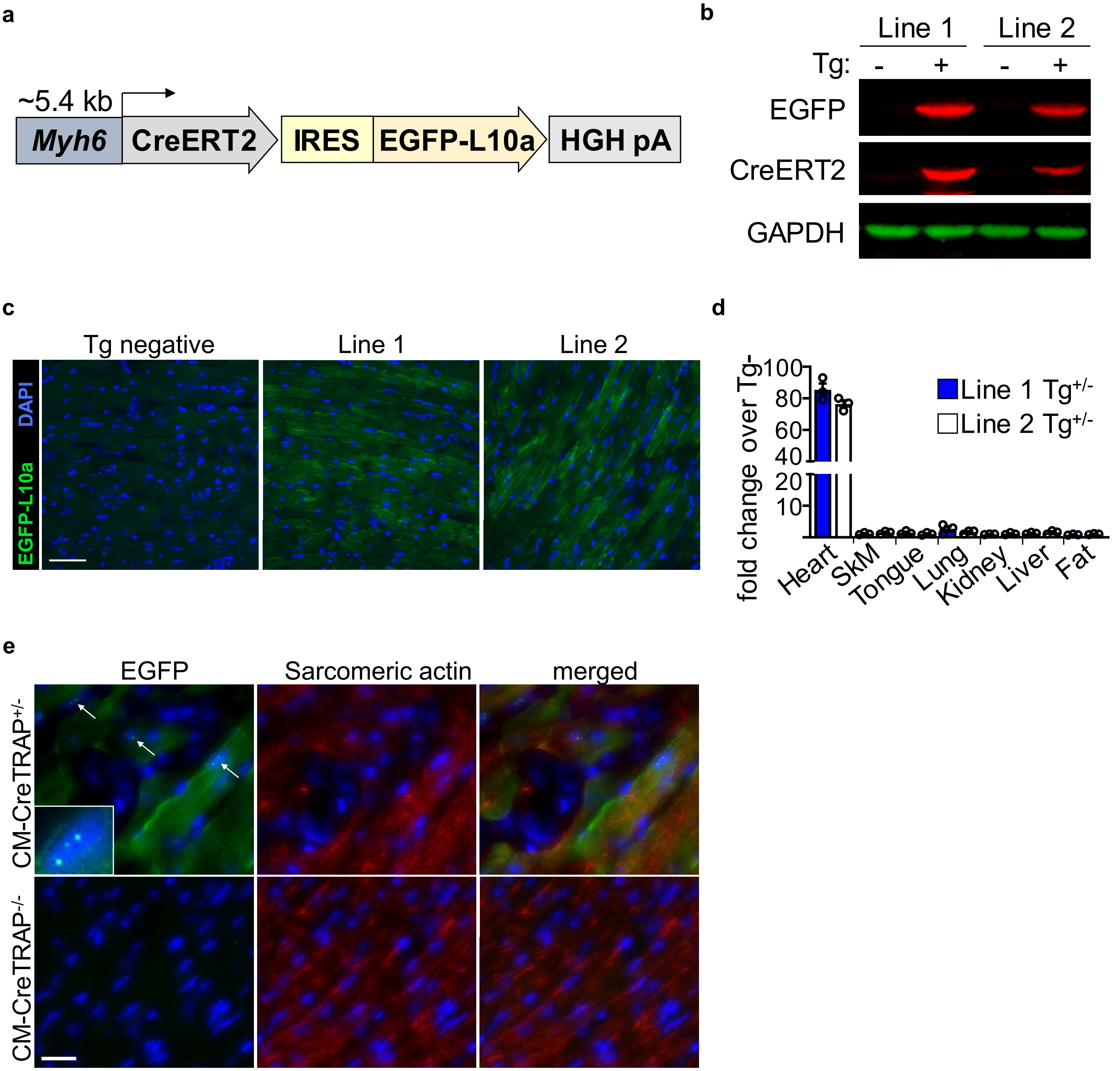


**Supplementary Fig. 1. Generation and EGFP-L10a expression in CM-CreTRAP mice.** (**a**) Schematic of the transgene vector. IRES and HGH pA denote internal ribosomal entry site and human growth hormone polyadenylation signal, respectively. (**b**) Immunoblot of GFP, CreERT2, and GAPDH on the heart extracts from 8-week-old lines 1 and 2 at F2. - and + signs denote transgenic (Tg) negative and positive, respectively. (**c**) Direct fluorescence micrograph images of heart sections from CM-CreTRAP transgenic (Tg) line 1 and line 2 showing robust green fluorescence signal but not in hearts from the Tg negative mice. DAPI counterstain shows the nucleus. Scale bar = 100µm. (**d**) qPCR of transgene (CreERT) mRNA expression (as fold change over Tg-) in various tissues normalized to Rpl13a from 8-week-old line 1 and 2 mice. SkM denotes skeletal muscle (quadriceps). Error bars indicate SEM. n = 3. (**e**) EGFP and sarcomeric actin staining on 8-week-old Tg+ and Tg- hearts from line 1. White arrows and inset highlight enriched areas of EGFP-L10a in the nucleolus. Scale bar = 30µm.

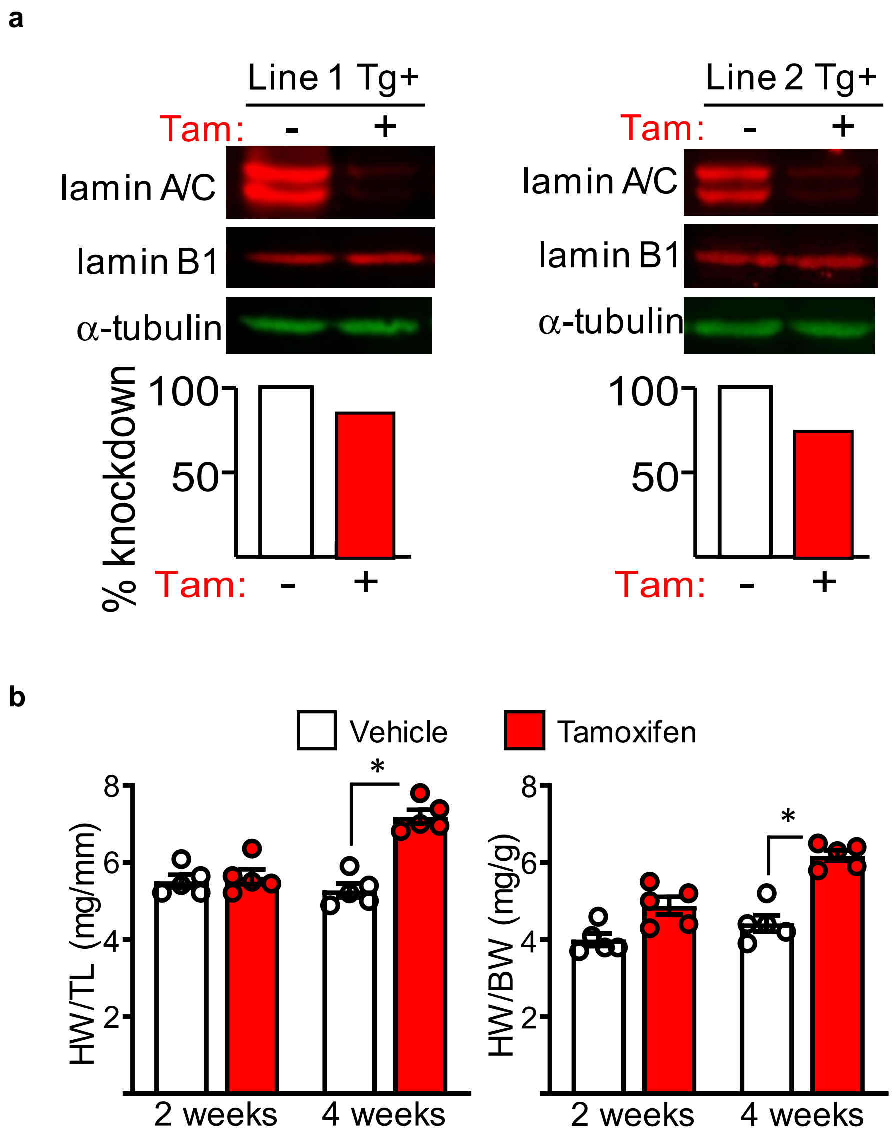


**Supplementary Fig. 2. CM-specific depletion of lamin A/C.** (**a**) Immunoblot of lamin A/C, lamin B1, and α-tubulin in CM extracts from Tg+ line 1 (left) and line 2 (right) mice (CM-CreTRAP:*Lmna*$^{flox/flox}$) with (+) or without (-) tamoxifen (Tam) administration. The bottom shows quantitation of knockdown. (**b**) Assessment of heart weight (HW) of CM-CreTRAP:*Lmna*$^{flox/flox}$ mice at 2 and 4 weeks post last Tam dosing. The left panel shows the HW relative to tibia length (TL) and the right panel relative to body weight (BW). P values were derived using unpaired, two tailed Student's t test. * = p < 0.0001. Error bars = SEM.

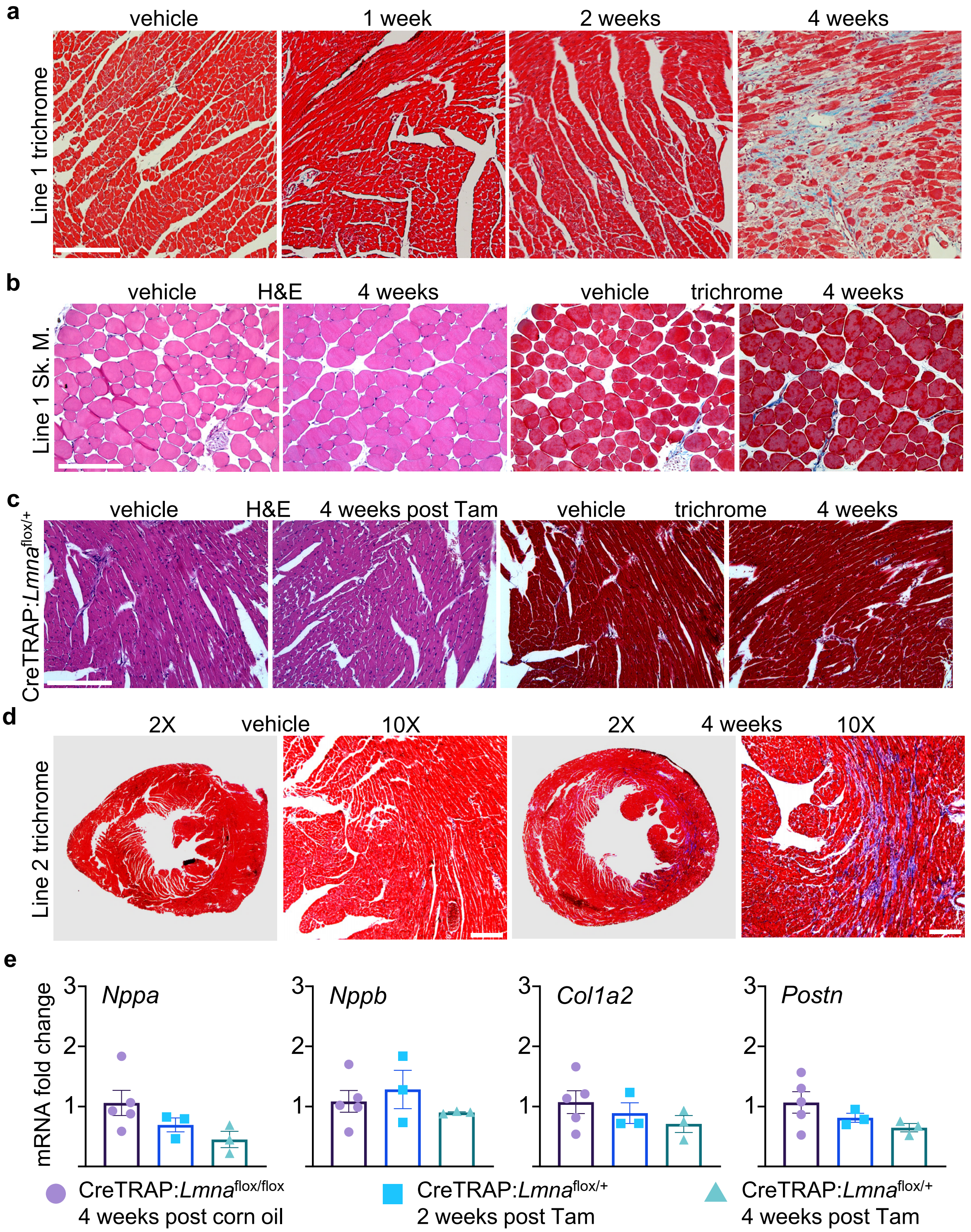


**Supplementary Fig. 3. CM-specific depletion of lamin A/C causes cardiac remodeling.** (**a**) Masson's trichrome staining of hearts from line 1 CM-CreTRAP:*Lmna*$^{flox/flox}$ mice at 1, 2, and 4 weeks post tamoxifen and vehicle treatment. (**b**) Hematoxylin and Eosin (H&E) and Masson's trichrome staining of quadriceps from line 1 CM-CreTRAP:*Lmna*$^{flox/flox}$ mice at 4 weeks post tamoxifen and vehicle treatment. Sk.M. denotes skeletal muscle. (**c**) H&E and Masson's trichrome staining of hearts from line 1 CM-CreTRAP:*Lmna*$^{flox/+}$ mice at 4 weeks post tamoxifen treatment. (**d**) Masson's trichrome staining of hearts from line 2 CM-CreTRAP:*Lmna*$^{flox/flox}$ mice at 4 weeks post tamoxifen and vehicle treatment. Representative images are shown from n =5 mice per group. Scale bars = 200µm. (**e**) qPCR of cardiac stress and profibrotic marker mRNAs in hearts from vehicle-treated CM-CreTRAP:*Lmna*$^{flox/flox}$ and Tam-treated CM-CreTRAP:*Lmna*$^{flox/+}$ mice. Data normalized to Rpl13a presented as fold change relative to vehicle (Veh). Error bars = SEM. n = 3 - 5.

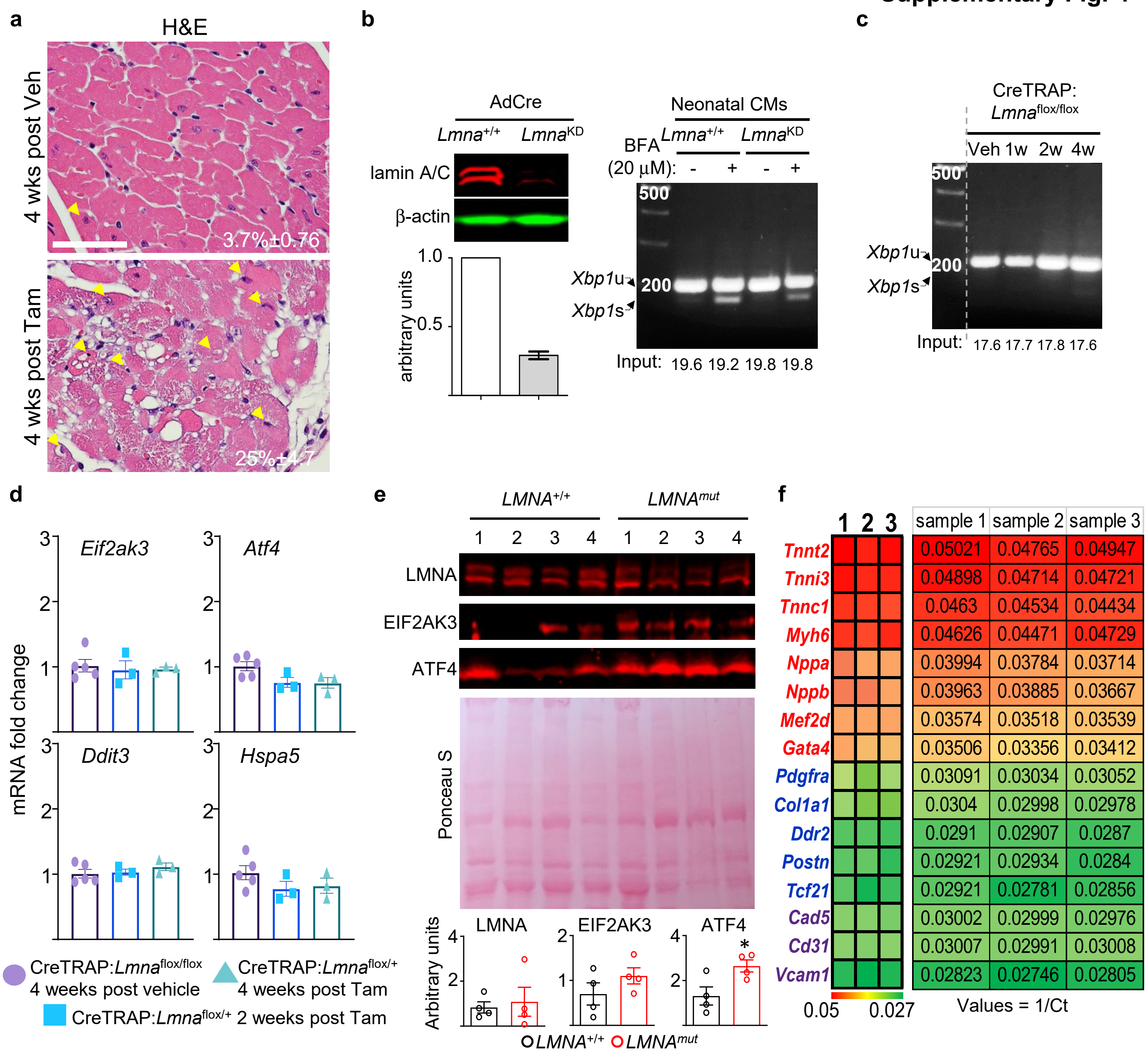


**Supplementary Fig. 4. Cardiac perturbations in CM-specific *Lmna* deleted mice.** (**a**) H&E staining of hearts from line 1 CM-CreTRAP:*Lmna*$^{flox/flox}$ mice at 4 weeks post tamoxifen and vehicle treatment. Yellow arrows denote abnormal nuclei. Representative images are shown. Scale bar = 50μm. (**b**) (Left panel) Immunoblot of lamin A/C and β-actin in AdCre-treated nCMs from *Lmna*$^{+/+}$ and *Lmna*$^{flox/flox}$ (*Lmna*$^{KD}$) mice. Quantitation of blots are shown on the bottom. (Right panel) RT-PCR analyses of *Xbp1* mRNA splicing in 24 hr brefeldin A (BFA)-treated *Lmna*$^{+/+}$ and *Lmna*-deleted neonatal CMs. *Xbp1*u and *Xbp1*s denote unspliced and spliced variants, respectively. Input denotes CT values for internal control *Rpl13a*. Representative images from 3 independent experiments are shown. (**c**) RT-PCR analyses of *Xbp1* mRNA splicing in the hearts from CM-CreTRAP:*Lmna*$^{flox/flox}$ mice treated with vehicle (Veh) or Tam, similar to those shown in Supplementary Fig. 4b. Input denotes CT values for internal control *Gapdh*. (**d**) qPCR of unfolded protein response marker mRNAs in hearts from vehicle-treated CM-CreTRAP:*Lmna*$^{flox/flox}$ and Tam-treated CM-CreTRAP:*Lmna*$^{flox/+}$ mice at 2 and 4 weeks post tamoxifen treatment. Data normalized to *Rpl13a* presented as fold change relative to vehicle (Veh). Error bars = SEM. n = 3 - 5. (**e**) Immunoblot of lamin A/C, PERK, and ATF4 on human hearts with quantitation on the bottom. Numbers on top of blots denote individual heart samples. Ponceau S stain was used to assess even loading. * denotes p = 0.03 using unpaired, two-tailed Student's t test. Error bars = SEM. (**f**) qPCR data on TRAP mRNA (n=3) in the hearts from CM-CreTRAP:*Lmna*$^{flox/flox}$ mice 2 weeks post vehicle or Tam treatment. Primers recognizing genes specifically expressed in CMs (red), cardiac fibroblasts (blue), and endothelial cells (purple) were used and presented as 1/cycle threshold (Ct) for each genes from 100ng TRAP mRNA. The actual values are shown on the right panel.

**a**

Number of reads

| Sample | Total | Aligned | % |
|---|---|---|---|
| veh.1 | 37,873,693 | 36,782,447 | 97% |
| veh.2 | 36,609,358 | 35,504,303 | 97% |
| veh.3 | 40,852,376 | 38,734,857 | 95% |
| 2w.1 | 37,500,006 | 36,235,769 | 97% |
| 2w.2 | 41,127,589 | 39,725,364 | 97% |
| 2w.3 | 33,520,115 | 32,338,371 | 96% |

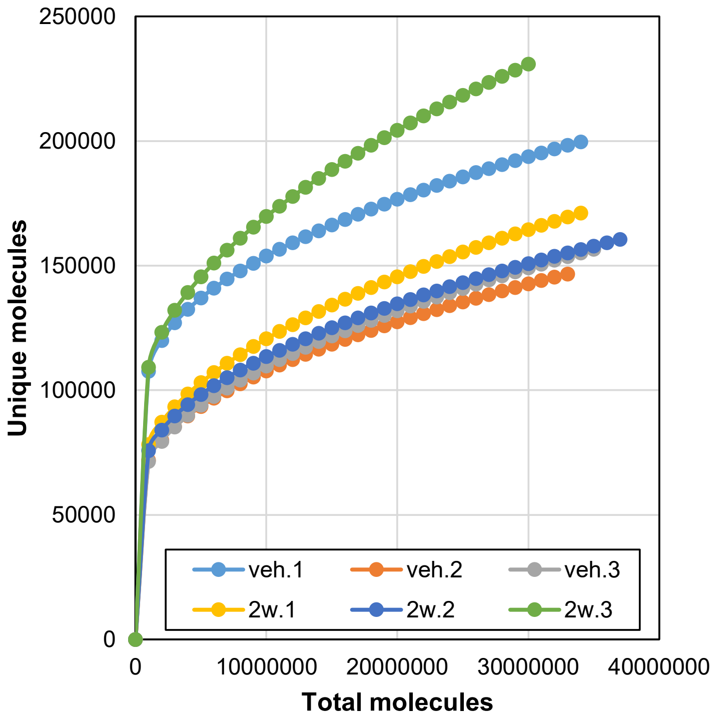


**b**

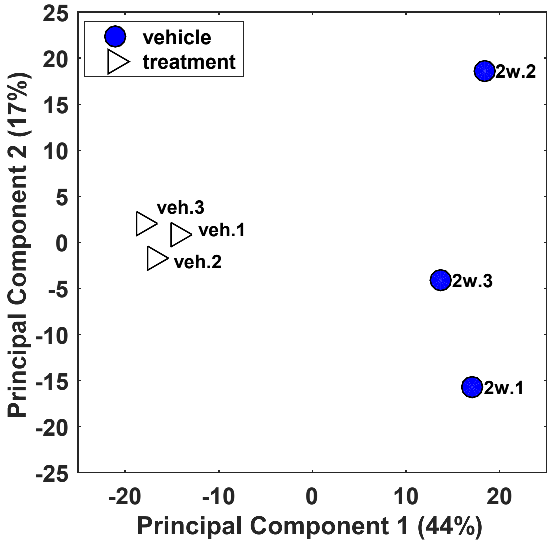


**c**

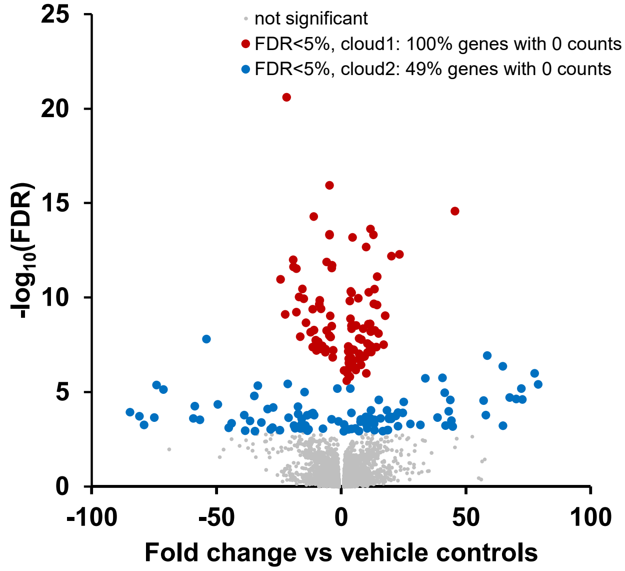


**d**

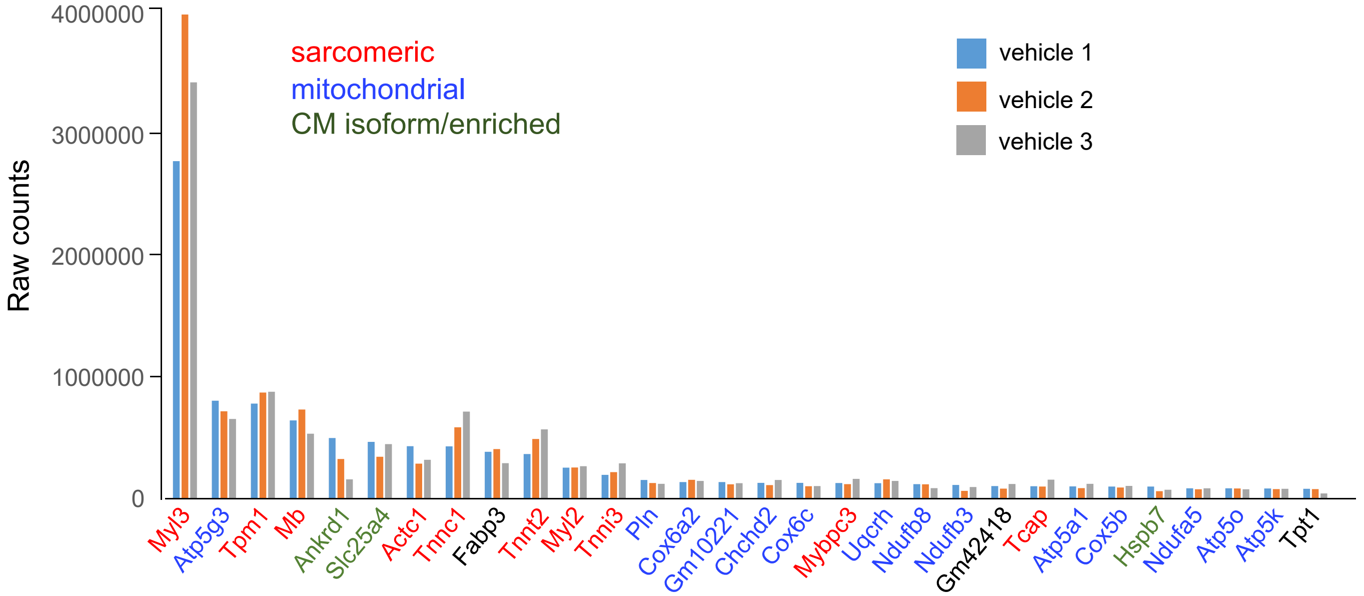


**Supplementary Fig. 5. Quality control data for TRAP sequencing.** (**a**) Top - Number of reads for TRAP sequencing. Bottom - unique molecule curve associated with the unique reads. Veh.1-3 and 2w.1-3 denote triplicates of TRAP sequencing samples from individual hearts. (**b**) Principle component analysis for the TRAP sequencing samples. (**c**) Volcano plot partitioned by genes with 100% zero counts. (**d**) Top 30 genes sorted based on raw counts from TRAP sequencing of hearts from vehicle treated mice (vehicle 1). Gene symbols with red fonts denote those encoding sarcomeric proteins, blue for mitochondrial, and green for CM-specific isoform or CM enriched.

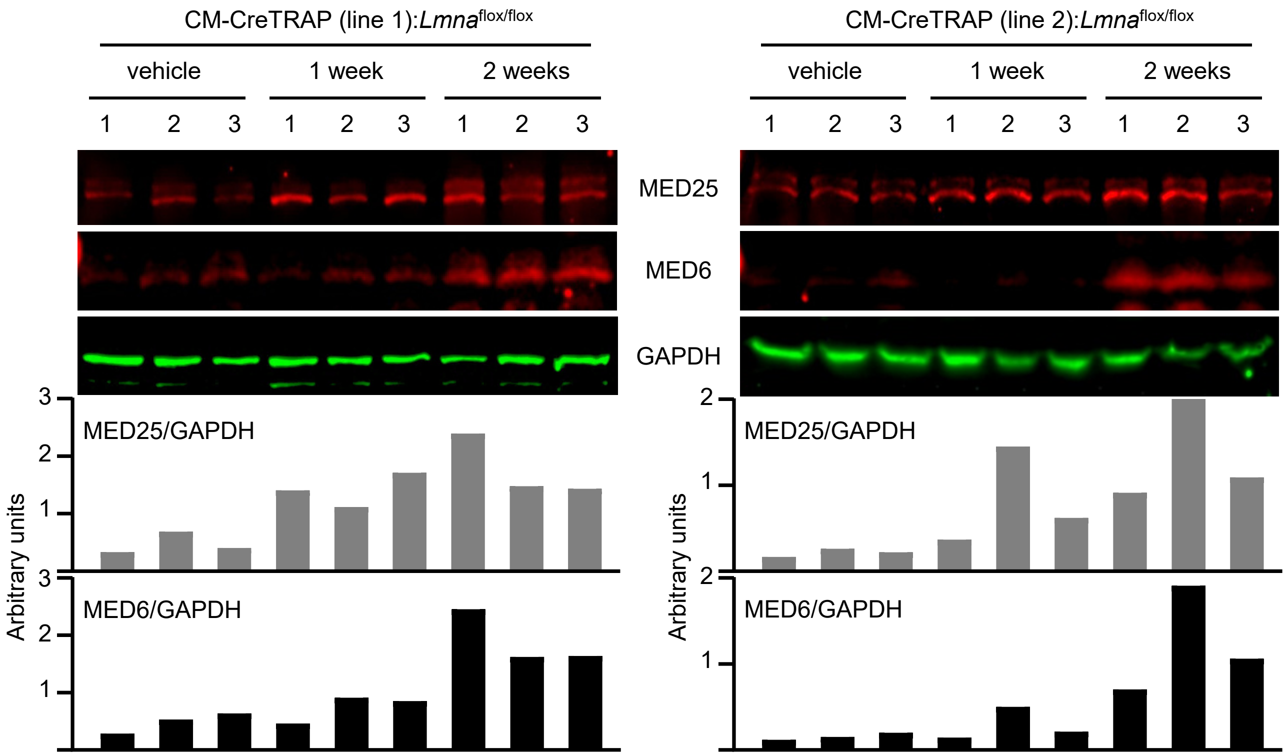


**Supplementary Fig. 6. Validation of MED25 and MED6 expression in *Lmna*-deleted hearts.** Immunoblot analysis of MED6, MED25, and GAPDH on the heart extracts from line 1 (left) and line 2 (right) CM-CreTRAP:*Lmna*$^{flox/flox}$ mice at 1 and 2 weeks post Tam treatment or vehicle alone (4W Tam). Numbers on top of blots denote individual heart samples. Bottom panel shows quantitation of blots normalized to GAPDH.

a

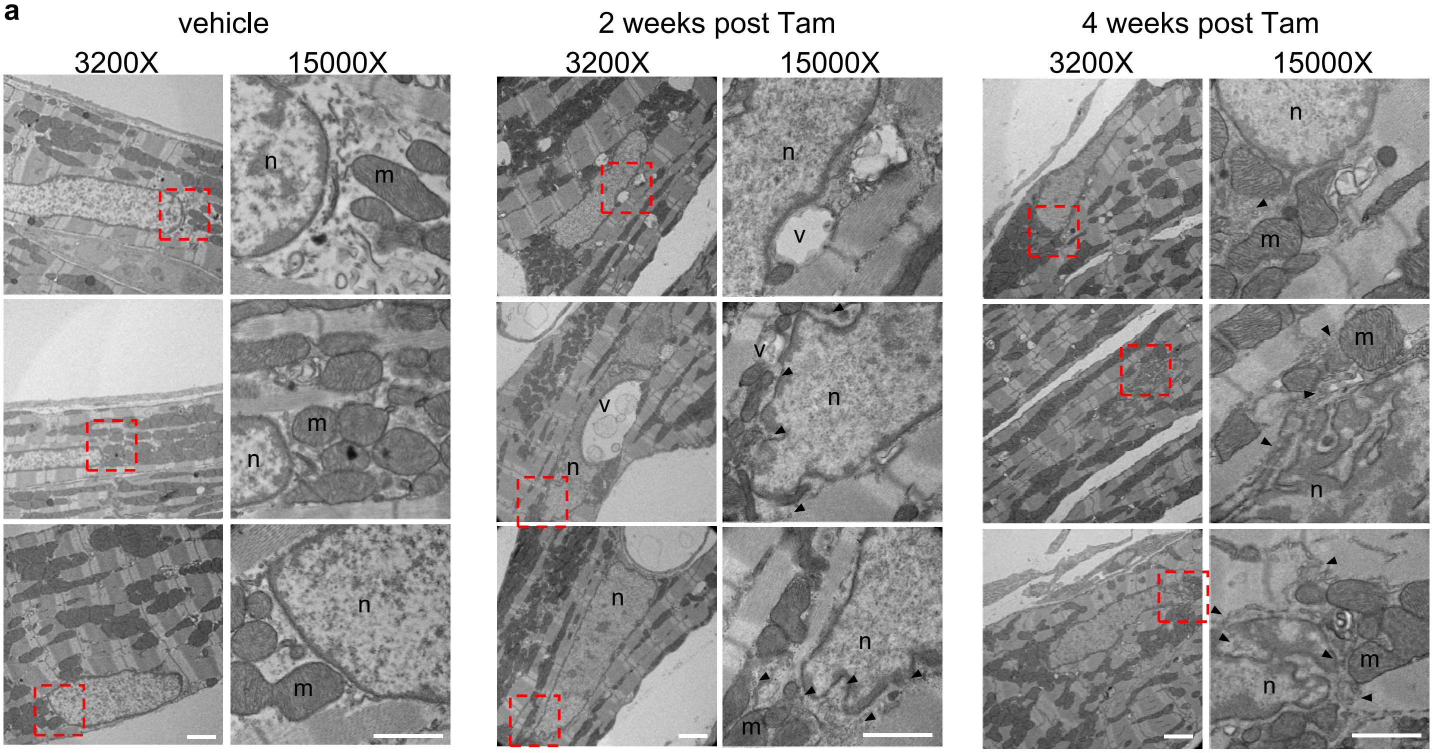


**Supplementary Fig. 7. Perinuclear abnormalities in hearts of CM-specific *Lmna* deleted mice.** (**a**) Transmission electron microscopy images of hearts from line 1 CM-CreTRAP:*Lmna*$^{flox/flox}$ mice treated with vehicle, 2 weeks, and 4 weeks post tamoxifen treatment. Dotted box denotes the area shown in the higher magnification (15000X) image. Scale bar = 1 µm. Black arrowheads denote 100 nm vesicles. n = nucleus, m = mitochondria, v = vacuoles. (**b**) Quantitation of EM images showing % of normal and abnormal nuclei from their respective groups. N = total number of nuclei counted from 2 independent hearts per group.

b

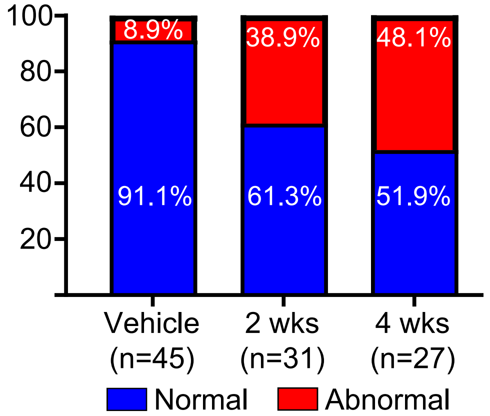

**Supplementary Fig. 8**

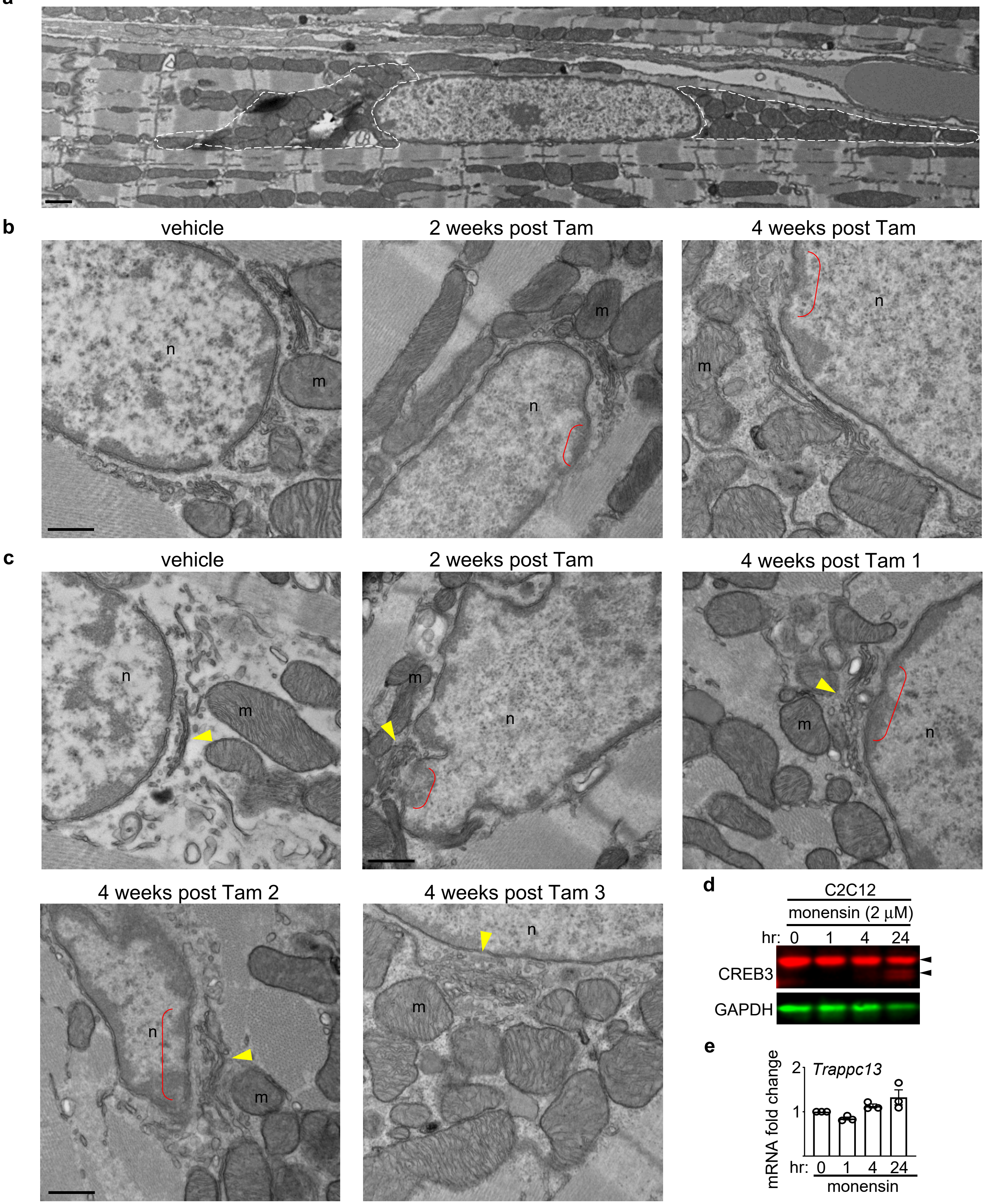


**Supplementary Fig. 8. Golgi abnormalities in hearts of CM-specific *Lmna* deleted mice.** (**a**) EM image of an adult CM showing perinuclear mitochondria highlighted by dashed lines. All scale bar in this figure = 500 nm. (**b**) Uncropped images of transmission electron microscopy images of hearts from line 1 CM-CreTRAP:*Lmna*$^{flox/flox}$ mice treated with vehicle, 2 weeks, and 4 weeks post tamoxifen treatment from fig. 4a. Red brackets show area of nuclear envelope deterioration. n = nucleus, m = mitochondria. (**c**) Additional EM images showing abnormal golgi (yellow arrowheads) in hearts from CM-CreTRAP:*Lmna*$^{flox/flox}$ mice treated with vehicle, 2 weeks, and 4 weeks post tamoxifen. Red brackets show area of nuclear envelope deterioration. (**d**) Immunoblot of CREB3 and GAPDH in C2C12 cells treated with 2 μM monensin for 1, 4, and 24 hrs. 0 hr denotes vehicle treatment (EtOH). Black arrowheads denote CREB3 and its fragments. (**e**) qPCR analysis of MEFs treated with 2 μM monensin for 0, 1, 4, and 24 hrs probed for *Trappc13*. n=3. Error bars denote SEM.

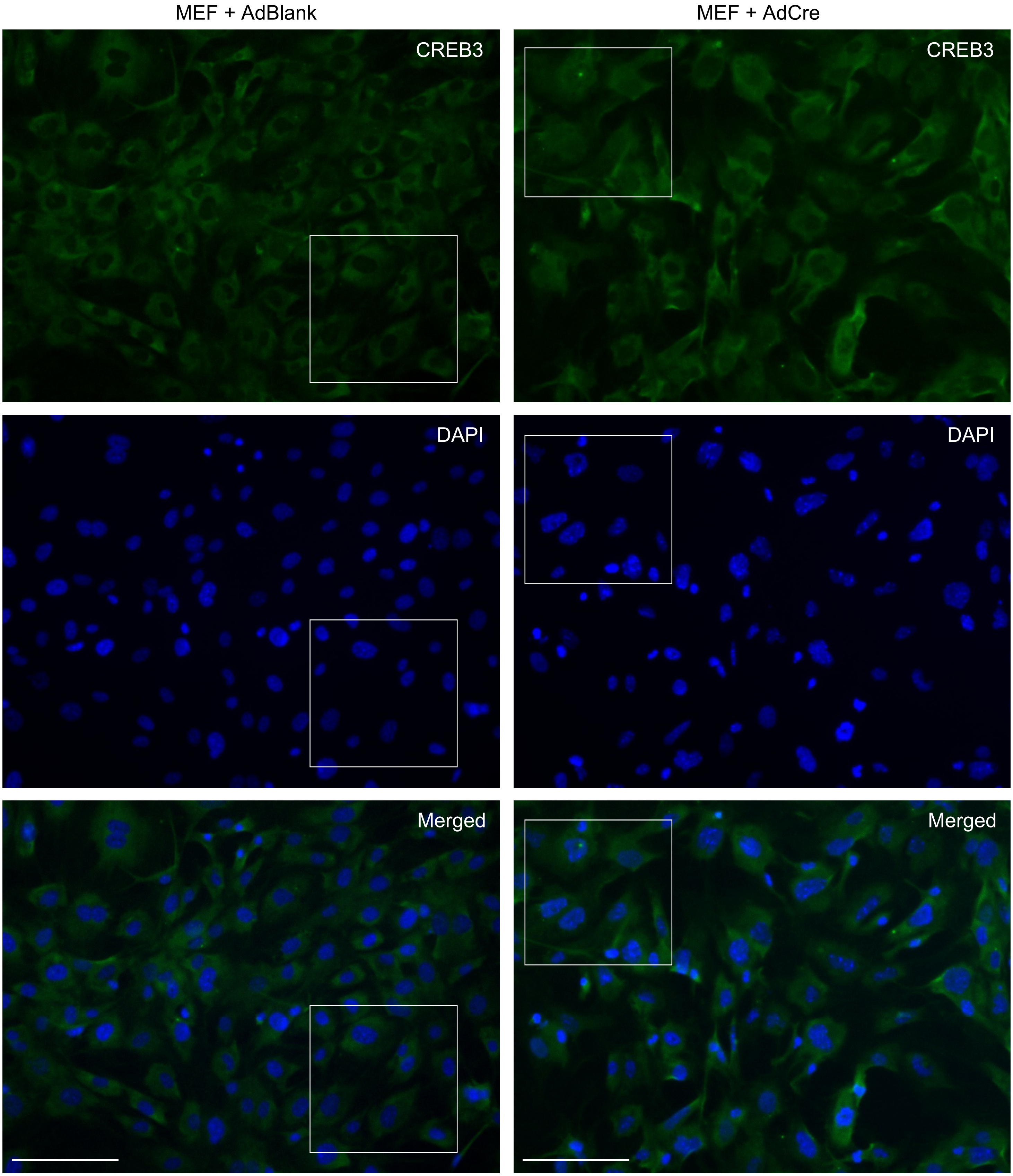


**Supplementary Fig. 9. Uncropped images shown in Figure 4e.** White boxes denote the region shown in Fig. 4e. Scale bar = 100 μm

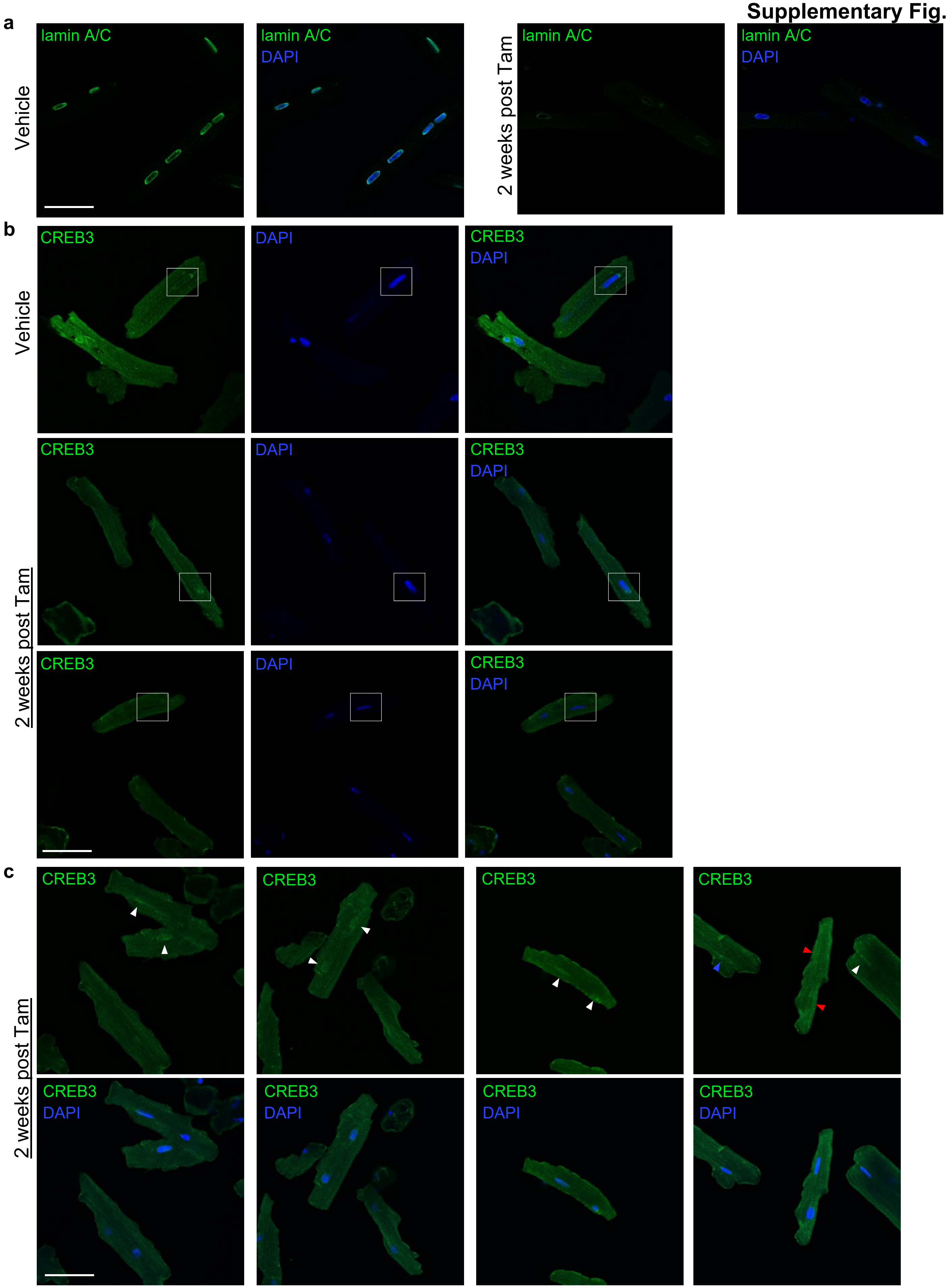


**Supplementary Fig. 10. CREB3 localization to the nucleus in adult CMs with *Lmna* deletion.** (**a**) Immunofluorescence images of adult CMs from hearts of CM-CreTRAP mice treated with Tam stained for lamin A/C and DAPI. (**b**) Uncropped images of adult CMs from hearts of vehicle or Tam-treated CM-CreTRAP mice stained for CREB3 and DAPI as shown in fig. 4f. (**c**) Additional uncropped images of adult CMs from hearts of Tam-treated CM-CreTRAP mice stained for CREB3 and DAPI. Scale bar = 50µm. Blue, red, and white arrowheads denote perinuclear, undetected, and intranuclear CREB3, respectively.

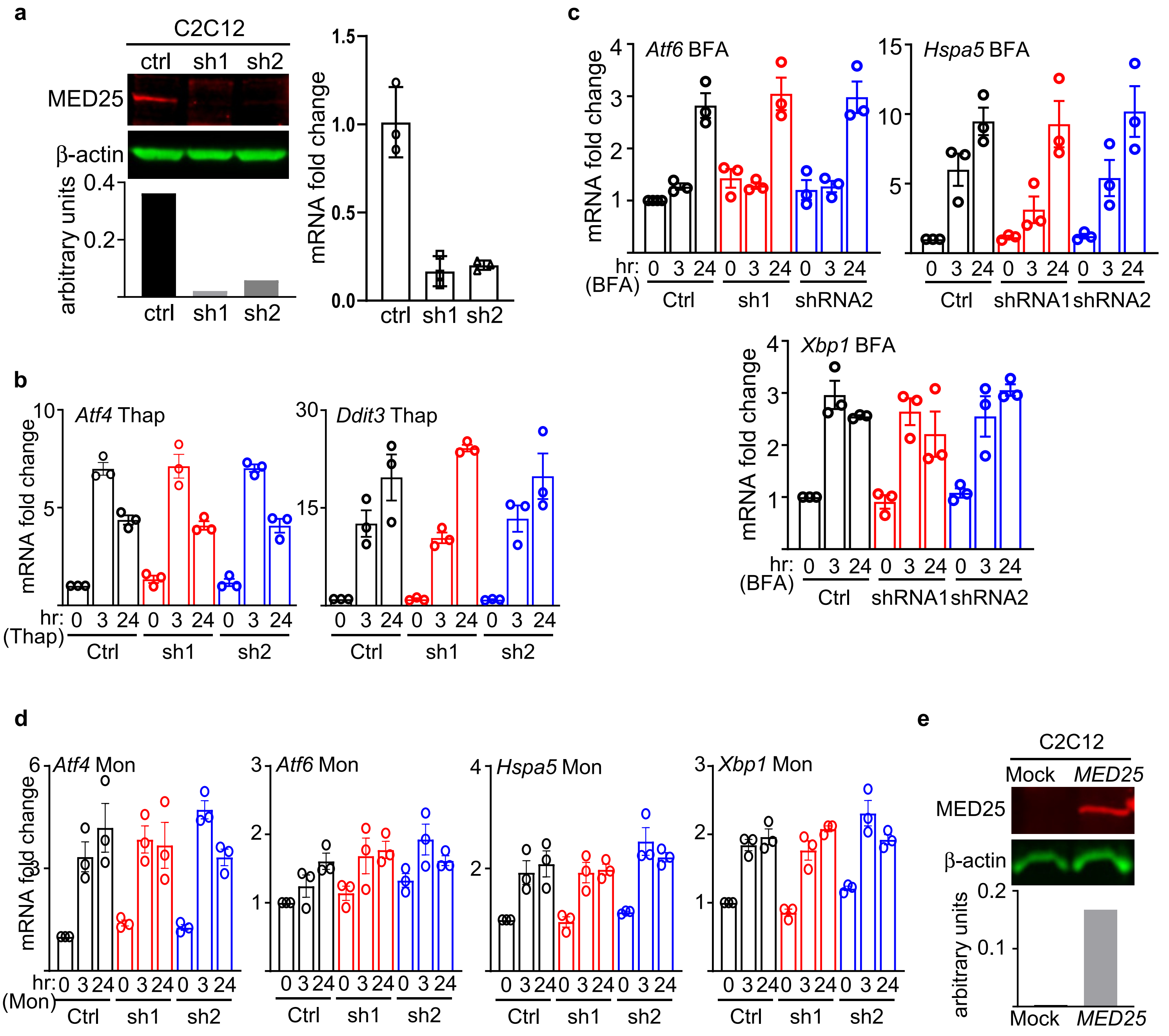


**Supplementary Fig. 11. *Med25* function in C2C12 cells.** (**a**) (Left panel) Immunoblot of MED25 and β-actin in nuclear extracts from C2C12 cells expressing two independent shRNAs (sh1 and sh2) that target *Med25*. Bottom panel shows quantitation of MED25 levels normalized to β-actin. (Right panel) qPCR analysis confirming *Med25* expression knockdown at the mRNA level. (**b**) qPCR analyses probing *Atf4* and *Ddit3* mRNA in MED25-depleted C2C12 cells treated with 2 µM thapsigargin for 3 and 24 hr. Fold change values were derived by setting untreated control as 1. (**c**) qPCR analyses probing *Atf6, Hspa5,* and *Xbp1* mRNA in MED25-depleted C2C12 cells treated with 20 µM brefeldin-A (BFA) for 3 and 24 hr. Fold change values were derived by setting untreated control as 1. (**d**) qPCR analyses probing *Atf4*, Atf6, Hspa5, and *Xbp1* mRNA in MED25-depleted C2C12 cells treated with 2 µM monensin (Mon) for 3 and 24 hr. Fold change values were derived in reference to untreated control (Ctrl) that were set as 1. For all qPCR data in this figure; n = 3 independent experiments. * = $p<0.05$. ** = $p<0.001$. (**e**) Immunoblot of MED25 (using FLAG antibodies) and β-actin in nuclear extracts from C2C12 cells transfected with FLAG-tagged *MED25* expression constructs. Bottom panel shows quantitation of MED25 normalized to β-actin. Representative data from 3 independent experiments are shown.

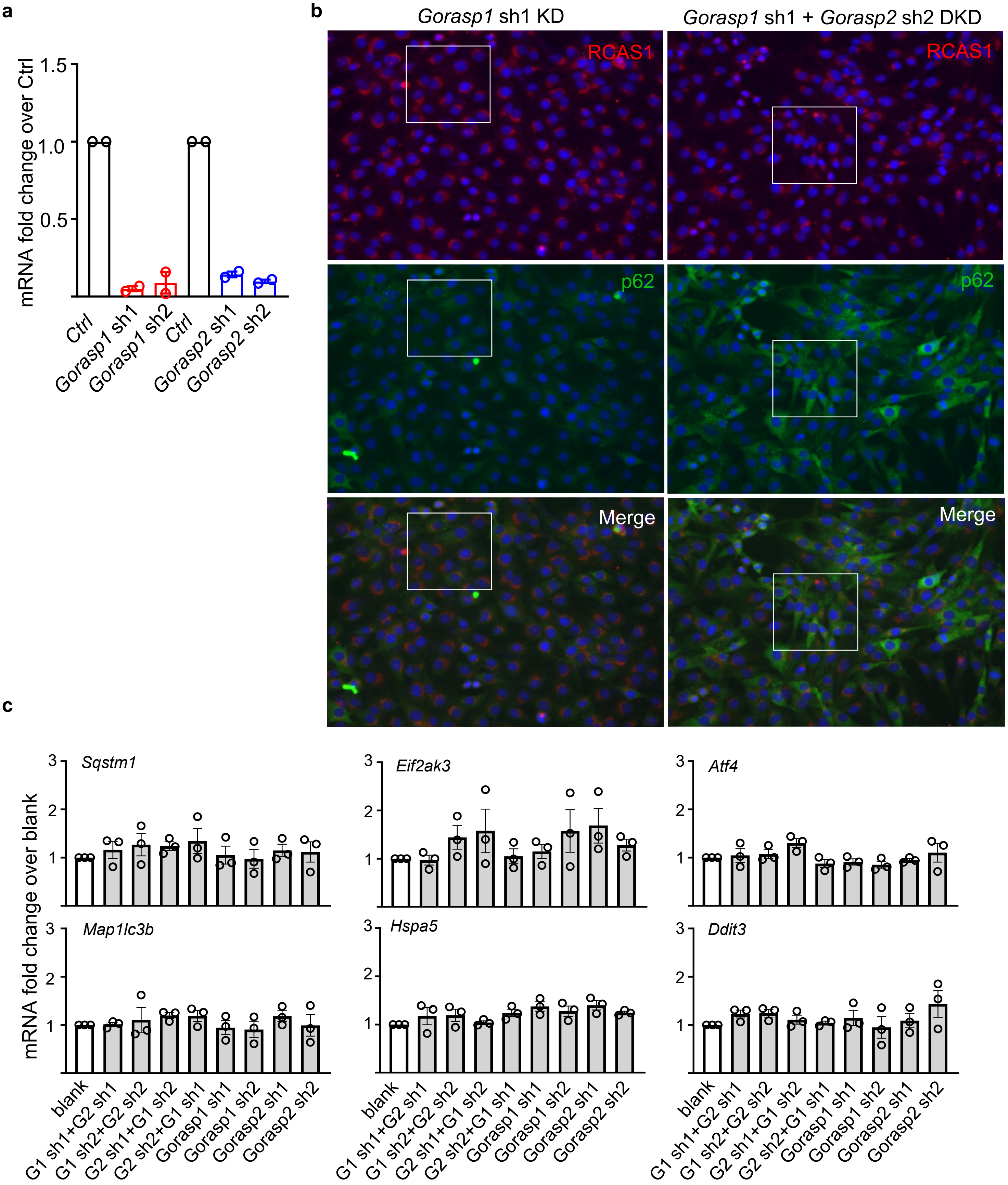


**Supplementary Fig. 12. Golgi disruption in C2C12 culture model.** (**a**) qPCR analyses of *Gorasp1* and *Gorasp2* knockdown in C2C12 using lentiviruses to deliver two independent shRNAs for each gene. Fold change values were derived using Ctrl, which is C2C12 infected with lentivirus carrying blank shRNA, as reference. Error bars = SEM. n = 2. (**b**) Uncropped images shown in Fig. 6c. Boxes show the cropped borders. Scale bar = 100 µm. (**c**) qPCR analyses on C2C12 cells with *Gorasp1* and *Gorasp2* knockdown probed for genes involved in UPR (*Eif2ak3*, *Atf4, Ddit3,* and *Hspa5*) and autophagy (*Sqstm1* and *Map1lc3b*). Error bars = SEM. n = 3.

a

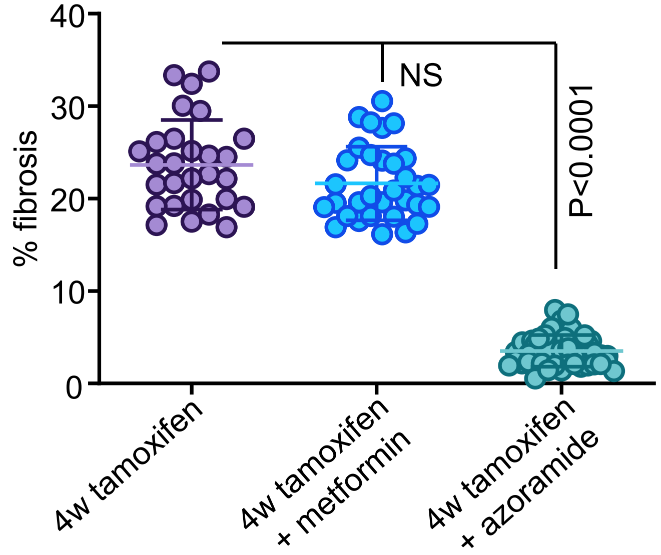


b

| | 4W Tam | | | | | 4W Tam + Metformin | | | | | | | | 4W Tam | | | | | 4W Tam+ Azoramide | | | | | | | |
|---|---|---|---|---|---|---|---|---|---|---|---|---|---|---|---|---|---|---|---|---|---|---|---|---|---|---|
| | **1** | **2** | **3** | **4** | **5** | **1** | **2** | **3** | **4** | **5** | **6** | **7** | **8** | **1** | **2** | **3** | **4** | **5** | **1** | **2** | **3** | **4** | **5** | **6** | **7** | **8** |
| *Nppa* | 0.5 | 1.4 | 0.9 | 1.7 | 1.0 | 1.3 | 1.2 | 1.2 | 1.3 | 1.0 | 1.9 | 1.3 | 0.9 | 0.9 | 1.1 | 0.7 | 1.5 | 1.0 | 0.4 | 0.4 | 0.2 | 0.1 | 0.3 | 0.6 | 0.5 | 0.2 |
| *Nppb* | 1.1 | 0.7 | 1.8 | 0.8 | 0.9 | 2.5 | 2.4 | 2.0 | 2.3 | 1.9 | 2.4 | 2.1 | 2.3 | 1.1 | 1.0 | 0.5 | 1.3 | 0.7 | 2.1 | 1.2 | 1.1 | 0.7 | 1.4 | 1.2 | 1.3 | 0.6 |
| *Acta2* | 1.0 | 0.5 | 1.6 | 0.9 | 1.3 | 1.1 | 3.1 | 0.9 | 1.4 | 3.9 | 1.7 | 2.1 | 3.3 | 1.0 | 0.6 | 1.4 | 1.2 | 1.2 | 1.0 | 1.1 | 1.0 | 0.9 | 1.0 | 0.9 | 1.3 | 0.7 |
| *Col1a1* | 1.8 | 0.6 | 1.6 | 1.4 | 0.4 | 1.7 | 1.0 | 1.6 | 1.3 | 1.1 | 1.7 | 1.7 | 1.2 | 1.0 | 1.9 | 0.5 | 0.9 | 1.2 | 0.8 | 0.6 | 0.6 | 0.2 | 0.6 | 0.8 | 0.4 | 0.4 |
| *Col1a2* | 1.5 | 0.8 | 1.1 | 1.5 | 0.5 | 1.5 | 0.8 | 1.5 | 1.0 | 0.9 | 1.5 | 1.6 | 1.1 | 1.4 | 0.6 | 1.0 | 0.9 | 1.4 | 0.8 | 0.6 | 0.5 | 0.2 | 0.6 | 0.7 | 0.4 | 0.4 |
| *Col3a2* | 1.9 | 0.5 | 1.6 | 1.4 | 0.5 | 2.6 | 1.6 | 2.1 | 1.7 | 2.2 | 2.5 | 2.6 | 1.6 | 1.6 | 0.9 | 1.0 | 1.1 | 1.0 | 1.1 | 1.0 | 0.8 | 0.3 | 0.8 | 1.1 | 0.6 | 0.5 |
| *Ctgf* | 0.3 | 1.6 | 2.4 | 1.7 | 0.6 | 1.3 | 2.3 | 3.4 | 3.4 | 1.6 | 4.1 | 3.5 | 4.4 | 1.0 | 1.0 | 0.9 | 1.0 | 1.3 | 0.8 | 0.6 | 0.4 | 0.3 | 0.6 | 0.5 | 0.5 | 0.4 |
| *Fn1-eda* | 1.5 | 0.2 | 2.0 | 2.3 | 0.8 | 1.7 | 0.4 | 1.1 | 0.8 | 0.9 | 2.5 | 0.8 | 0.1 | 1.5 | 0.8 | 0.8 | 1.1 | 1.1 | 1.8 | 1.6 | 1.1 | 0.1 | 0.5 | 1.6 | 0.8 | 1.4 |
| *Postn* | 1.7 | 0.6 | 1.5 | 1.4 | 0.5 | 1.2 | 0.7 | 0.9 | 0.7 | 1.0 | 1.4 | 1.1 | 0.9 | 1.3 | 1.1 | 1.3 | 0.6 | 1.2 | 0.6 | 0.5 | 0.4 | 0.1 | 0.4 | 0.4 | 0.3 | 0.1 |
| *Atf4* | 1.2 | 1.2 | 0.9 | 0.9 | 0.9 | 0.9 | 1.0 | 0.9 | 0.8 | 0.8 | 0.9 | 1.0 | 1.2 | 1.1 | 1.0 | 0.9 | 1.0 | 1.3 | 0.9 | 0.7 | 0.7 | 0.8 | 0.8 | 0.9 | 0.7 | 0.8 |
| *Ddit3* | 1.0 | 1.0 | 0.8 | 1.1 | 1.2 | 1.0 | 1.1 | 0.9 | 0.7 | 1.0 | 0.7 | 1.0 | 0.9 | 1.0 | 1.1 | 1.0 | 0.7 | 1.2 | 0.5 | 0.6 | 0.5 | 0.5 | 0.6 | 0.6 | 0.6 | 0.6 |
| *Eif2ak3* | 0.9 | 0.9 | 1.2 | 1.2 | 1.0 | 1.1 | 1.1 | 1.1 | 1.1 | 1.4 | 1.2 | 1.4 | 1.4 | 1.6 | 0.8 | 0.8 | 1.0 | 1.3 | 0.7 | 0.7 | 0.7 | 0.7 | 0.7 | 0.8 | 0.6 | 0.7 |
| *Hspa5* | 0.9 | 1.1 | 0.8 | 1.2 | 1.1 | 1.0 | 1.0 | 0.8 | 1.0 | 1.3 | 1.0 | 1.1 | 1.2 | 1.1 | 1.5 | 0.8 | 1.0 | 1.2 | 0.6 | 0.6 | 0.5 | 0.5 | 0.6 | 0.6 | 0.5 | 0.5 |
| *Atf6* | 0.8 | 1.2 | 0.9 | 1.1 | 0.7 | 1.0 | 1.2 | 1.0 | 1.0 | 1.2 | 1.4 | 1.1 | 0.9 | 0.9 | 1.1 | 0.9 | 1.1 | 0.9 | 1.0 | 1.3 | 0.9 | 1.2 | 1.0 | 1.0 | 0.9 | 1.2 |
| *Xbp1* | 1.3 | 0.9 | 1.0 | 1.0 | 1.0 | 1.0 | 1.1 | 1.4 | 1.2 | 1.6 | 1.0 | 1.4 | 1.1 | 1.4 | 0.8 | 1.0 | 1.0 | 1.1 | 1.0 | 1.0 | 0.9 | 1.0 | 1.0 | 1.1 | 0.8 | 1.0 |
| *Atg5* | 1.2 | 0.9 | 1.2 | 0.9 | 0.9 | 1.5 | 1.3 | 1.3 | 1.4 | 1.4 | 1.3 | 1.7 | 1.8 | 1.2 | 0.9 | 0.8 | 0.9 | 0.9 | 1.2 | 0.8 | 0.8 | 0.8 | 0.7 | 0.7 | 0.7 | 0.8 |
| *Atg7* | 1.1 | 1.0 | 1.1 | 1.0 | 0.9 | 1.3 | 1.1 | 1.7 | 1.1 | 1.4 | 1.1 | 1.6 | 1.7 | 1.2 | 1.2 | 1.0 | 1.1 | 0.9 | 0.9 | 1.0 | 0.8 | 0.9 | 0.8 | 1.1 | 0.9 | 1.1 |
| *Becn1* | 1.0 | 1.1 | 1.0 | 1.0 | 0.9 | 1.2 | 1.4 | 1.1 | 1.1 | 1.9 | 1.0 | 1.7 | 1.7 | 1.2 | 1.3 | 0.9 | 0.8 | 1.0 | 0.7 | 1.0 | 1.1 | 0.8 | 0.7 | 0.8 | 0.8 | 0.9 |
| *Gabarap* | 0.7 | 0.8 | 1.5 | 1.0 | 1.2 | 1.7 | 1.1 | 2.0 | 2.1 | 2.7 | 2.6 | 3.6 | 2.5 | 1.4 | 1.2 | 0.7 | 0.9 | 1.3 | 0.9 | 1.5 | 0.7 | 1.3 | 1.4 | 1.9 | 1.4 | 0.9 |
| *Gabarapl1* | 0.6 | 0.8 | 1.2 | 1.2 | 1.3 | 1.5 | 1.1 | 2.0 | 1.2 | 2.3 | 1.5 | 2.0 | 2.2 | 1.5 | 0.8 | 0.8 | 0.8 | 1.2 | 1.0 | 0.5 | 0.8 | 0.9 | 0.6 | 0.6 | 0.7 | 0.9 |
| *Lamp2* | 1.3 | 0.9 | 1.0 | 0.9 | 0.9 | 0.9 | 0.9 | 1.2 | 1.1 | 1.2 | 0.9 | 1.2 | 1.3 | 1.0 | 1.0 | 1.0 | 0.9 | 1.1 | 0.8 | 0.9 | 0.8 | 0.8 | 0.9 | 0.8 | 0.9 | 0.9 |
| *Map1lc3a* | 1.0 | 1.0 | 1.0 | 0.9 | 1.1 | 1.3 | 1.5 | 1.0 | 1.5 | 2.0 | 1.3 | 2.1 | 1.9 | 1.4 | 1.0 | 0.8 | 0.9 | 1.0 | 0.8 | 1.0 | 0.9 | 1.1 | 1.1 | 1.0 | 1.1 | 1.3 |
| *Map1lc3b* | 1.1 | 1.1 | 1.0 | 1.0 | 0.9 | 0.9 | 1.1 | 0.9 | 1.0 | 1.6 | 1.0 | 1.4 | 1.1 | 1.1 | 1.1 | 1.0 | 1.0 | 1.0 | 0.9 | 1.1 | 0.9 | 1.0 | 1.6 | 1.0 | 1.4 | 1.1 |
| *Pik3c3* | 1.0 | 0.9 | 1.0 | 1.2 | 0.9 | 1.0 | 1.1 | 1.4 | 1.2 | 1.6 | 1.0 | 1.7 | 1.6 | 1.0 | 0.9 | 0.9 | 1.2 | 0.9 | 0.8 | 1.0 | 1.0 | 0.9 | 0.9 | 0.9 | 1.0 | 1.0 |
| *Tfeb* | 1.6 | 0.5 | 1.5 | 1.0 | 0.9 | 1.8 | 2.1 | 2.4 | 2.2 | 3.0 | 2.1 | 3.2 | 3.2 | 1.4 | 0.7 | 0.8 | 0.9 | 1.3 | 1.6 | 1.8 | 1.8 | 1.8 | 1.5 | 2.1 | 1.9 | 2.2 |
| *Ulk1* | 1.0 | 1.2 | 1.1 | 0.9 | 0.8 | 1.0 | 0.8 | 1.3 | 1.1 | 1.4 | 1.1 | 1.7 | 1.7 | 1.3 | 1.0 | 1.6 | 0.7 | 0.8 | 0.7 | 0.8 | 0.6 | 1.0 | 1.1 | 1.1 | 1.3 | 1.3 |
| *Tnfrsf1b* | 1.2 | 1.1 | 0.9 | 1.0 | 0.9 | 0.7 | 0.8 | 1.2 | 0.7 | 0.8 | 1.2 | 0.7 | 0.9 | 1.0 | 0.9 | 0.9 | 1.4 | 1.0 | 0.3 | 0.8 | 0.7 | 0.5 | 0.6 | 0.7 | 0.8 | 1.0 |
| *Tnfrsf10b* | 1.2 | 1.1 | 0.9 | 1.1 | 0.8 | 0.7 | 1.1 | 1.4 | 1.0 | 0.9 | 0.7 | 0.9 | 0.9 | 1.0 | 1.0 | 1.2 | 1.0 | 0.8 | 0.8 | 0.9 | 1.0 | 0.3 | 0.4 | 0.4 | 0.5 | 1.0 |
| *Arf4* | 1.1 | 1.0 | 0.8 | 1.3 | 0.9 | 0.7 | 0.8 | 0.9 | 0.8 | 0.7 | 1.1 | 0.7 | 0.9 | 0.7 | 1.0 | 0.8 | 1.0 | 1.8 | 0.7 | 1.7 | 1.0 | 0.6 | 0.6 | 0.8 | 1.0 | 1.1 |

**Supplementary Fig. 13. Characterization of Tam-treated CM-CreTRAP:*Lmna*$^{flox/flox}$ mice treated with metformin or azoramide.** (**a**) Quantitation of % fibrosis based on Masson's trichrome staining of hearts from CM-CreTRAP:*Lmna*$^{flox/flox}$ mice at 4 weeks post Tam treatment alone or with either metformin or azoramide. Average % fibrosis was calculated from ~30 independent images from 3 mouse hearts per group. Error bars = standard deviation. NS = not significant. (**b**) mRNA fold change values obtained from qPCR analyses to generate the heatmap shown in fig. 7d. Numbers on top denote individual heart samples. Average ΔCt values from 4W Tam (n=5) were used as a reference to determine all ΔΔCt fold change values.

| Groups | Sex(n) | FS (%) | EF (%) | LVESD(mm) | LVEDD(mm) | LVESPW(mm) | LVEDPW(mm) |
|---|---|---|---|---|---|---|---|
| CreTRAP:*Lmna*$^{flox/flox}$ + corn oil (Veh) | M (5) | 27.4±0.49 | 53.4±0.79 | 3.22±0.076 | 4.44±0.096 | 1.1±0.041 | 0.78±0.042 |
| | F (5) | 28.4±1.85 | 54.7±2.89 | 3.18±0.076 | 4.43±0.085 | 1.08±0.057 | 0.9±0.075 |
| CreTRAP:*Lmna*$^{flox/+}$ + 2 weeks post Tam | M (5) | 26.7±0.51 | 52.4±0.90 | 3.19±0.117 | 4.34±0.130 | 1.15±0.029 | 0.82±0.017 |
| CreTRAP:*Lmna*$^{flox/+}$ + 4 weeks post Tam | M (5) | 25.3±1.15 | 49.9±1.85 | 3.32±0.091 | 4.45±0.114 | 1.08±0.035 | 0.78±0.056 |
| CreTRAP:*Lmna*$^{flox/flox}$ + 2 weeks post Tam | M (5) | 24.5±1.08 | 48.7±1.78 | 3.45±0.068 | 4.57±0.074 | 1.28±0.053 | 0.93±0.079 |
| | F (5) | 28.8±3.55 | 55.4±5.74 | 2.80±0.296 | 3.88±0.215 | 1.23±0.11 | 0.96±0.099 |
| CreTRAP:*Lmna*$^{flox/flox}$ + 4 weeks post Tam | M (6) | 10.1±3.23 | 22.4±6.40 | 3.81±0.256 | 4.24±0.177 | 0.99±0.159 | 0.87±0.12 |
| | F (4) | 9.91±2.93 | 21.3±5.60 | 3.72±0.160 | 4.13±0.062 | 0.89±0.089 | 0.8±0.038 |

**Supplementary Table 1. Echocardiography table for Fig. 1d.** FS = fractional shortening, EF = ejection fraction, LVESD = left ventricular end systolic dimension, LVEDD = left ventricular end diastolic dimension, LVESPW = left ventricular end systolic posterior wall thickness, LVEDPW = left ventricular end diastolic posterior wall thickness.

| Groups | Sex(n) | FS (%) | EF (%) | LVESD(mm) | LVEDD(mm) | LVESPW(mm) | LVEDPW(mm) |
|---|---|---|---|---|---|---|---|
| CreTRAP:*Lmna*$^{flox/flox}$ + corn oil (Veh) | M (7) / F (5) | 28.2±1.24 | 55.5±1.47 | 2.99±0.083 | 4.16±0.058 | 1.13±0.039 | 0.82±0.02 |
| CreTRAP:*Lmna*$^{flox/flox}$ + 2 weeks post Tam | M (6) / F (6) | 24.7±1.13 | 48.3±1.81 | 3.22±0.085 | 4.27±0.076 | 1.12±0.063 | 0.84±0.046 |
| CreTRAP:*Lmna*$^{flox/flox}$ + 2 weeks Tam+Met | M (3) / F (5) | 25.9±2.62 | 50.8±4.15 | 2.93±0.156 | 3.93±0.97 | 1.1±0.065 | 0.77±0.05 |
| CreTRAP:*Lmna*$^{flox/flox}$ + 2 weeks Tam+Azor | M (6) / F (2) | 28.3±2.09 | 54.9±3.23 | 2.79±0.144 | 3.89±0.132 | 1.07±0.05 | 0.71±0.038 |
| CreTRAP:*Lmna*$^{flox/flox}$ + 4 weeks post Tam | M (6) / F (6) | 9.02±1.43 | 19.6±2.99 | 4.29±0.153 | 4.7±0.114 | 0.893±0.05 | 0.85±0.053 |
| CreTRAP:*Lmna*$^{flox/flox}$ + 4 weeks Tam+Met | M (3) / F (5) | 19.3±1.91 | 39.7±3.44 | 3.43±0.201 | 4.23±0.165 | 1.1±0.07 | 0.86±0.059 |
| CreTRAP:*Lmna*$^{flox/flox}$ + 4 weeks Tam+Azor | M (6) / F (2) | 24.7±2.8 | 48.8±4.74 | 2.91±0.149 | 3.84±0.084 | 1.08±0.048 | 0.79±0.044 |

**Supplementary Table 2. Echocardiography table for Fig. 7b.** FS = fractional shortening, EF = ejection fraction, LVESD = left ventricular end systolic dimension, LVEDD = left ventricular end diastolic dimension, LVESPW = left ventricular end systolic posterior wall thickness, LVEDPW = left ventricular end diastolic posterior wall thickness.

Supplementary Table 3

| Primers | Forward (5' - 3') | Reverse (5' - 3') |
|---|---|---|
| ***CreERT*** | CGT ACT GAC GGT GGG AGA AT | CCC GGC AAA ACA GGT AGT TA |
| ***Nppa*** | TCG TCT TGG CCT TTT GGC T | TCC AGG TGG TCT AGC AGG TTC T |
| ***Nppb*** | AAG TCC TAG CCA GTC TCC AGA | GAG CTG TCT CTG GGC CAT TTC |
| ***Ccn2* (CTGF)** | GTG CCA GAA CGC ACA CTG | CCC CGG TTA CAC TCC AAA |
| ***Postn*** | ATG TCA TTG ACC GTG TCC TG | AAG AGC GTG AAG TGA CCA TC |
| ***Col1a1*** | TTC TCC TGG CAA AGA CGG ACT CAA | AGG AAG CTG AAG TCA TAA CCG CCA |
| ***Col1a2*** | GGC CCC CTG GTA TGA CTG GCT | CGC CAC GGG GAC CAC GAA TC |
| ***Col3a1*** | GTT CTA GAG GATGGCTGTACTAAACACA | TTG CCT TGC GTG TTT GAT ATT C |
| ***Fn1 EDA*** | CAG AAA TGA CCA TTG AAG GT | ATG AGT CCT GAC ACA ATC AC |
| ***Eif2ak3* (PERK)** | TCC CTG CTC GAA TCT TCC TA | CAT CCC AAG GCA GAA CAG AT |
| ***Atf4*** | ATG GGT TCT CCA GCG ACA | TCC ATT TTC TCC AAC ATC CAA |
| ***Ddit3* (CHOP)** | CTG GAA GCC TGG TAT GAG GA | CCT CTG TCA GCC AAG CTA GG |
| ***Hspa5* (BiP)** | CTT GGG GAC CAC CTA TTC CT | GGT TGG ACG TGA GTT GGT TC |
| ***Atf6*** | GCA GCA GTC GAT TAT CAG CA | GTT AGG TAG CTG TGC GGC TC |
| ***Xbp1* total** | TAT CCT TTT GGG CAT TCT GG | ACA GAG AAA GGG AGG CTG GT |
| ***Xbp1* spliced** | GAA CCA GGA GTT AAG AAC ACG | AGG CAA CAG TGT CAG AGT CC |
| ***Tnnt2*** | TAC AGA CTC TGA TCG AGG CTC ACT TC | TCA TTG CGA ATA CGC TGC TGC TC |
| ***Tnni3*** | TAA GAT CTC CGC CTC CAG AA | CGG CAT AAG TCC TGA AGC TC |
| ***Tnnc1*** | AGG CAG CCT TGA ACT CAT TC | TGT CCT GTG AGC TGT CTC CA |
| ***Myh6*** | ACG GTG ACC ATA AAG GAG GA | TGT CCT CGA TCT TGT CGA AC |
| ***Mef2d*** | TCA ACC ACT CCA ACA AGC TGT T | GTA CTC GGT GTA CTT GAG CAG CA |
| ***Gata4*** | GAG CCT GCC AAG CCA AGC | CTC CCG TCT ATC ACC TTT GTC C |
| ***Pdgfra*** | GGG AAG GAC TGG AAG CTT GGG GC | AGA TGA GGC CCG GCC CTG TGA GG |
| ***Ddr2*** | TTC CCT GCC CAG CGA GTC CA | ACC ACT GCA CCC TGA CTC CTC C |
| ***Tcf21*** | ATG CTG GAC TGT GAC TCC CT | GAG CGG GCT TTT CTT AGT GG |
| ***Cdh5*** | TCT TGC CAG CAA ACT CTC CT | TTG GAA TCA AAT GCA CAT CG |
| ***Pecam1*** | TCA CCA TCA ACA GCA TCC A | GGT GCT GAG ACC TGC TTT TC |
| ***Vcam1*** | CCG GCA TAT ACG AGT GTG AA | GAT GCG CAG TAG AGT GCA AG |
| ***Med25*** | GTG GTG GCG AGA GCT GTA GT | TCT CAA ACA GAA GCC GGA G |
| ***Atg5*** | ACA GCT GCA CAC ACT TGG AG | TTC CAG CAT TGG CTC TAT CC |
| ***Atg7*** | CCA GGA CAC CCT GTG AAC TT | GCT CTC CCT GGT GTC CAT TA |
| ***Becn1*** | CAG CTG GAC ACT CAG CTC AA | CTT GCG GTT CTT TTC CAC AT |
| ***Gabarap*** | TCC CGG TGA TAG TGG AAA AA | AAT TCG CTT CCG GAT CAA G |
| ***Gabarapl1*** | GAC CTC ACT GTT GGC CAG TT | TCT TCC TCG TGG TTG TCC TC |
| ***Lamp2*** | AGA CCA AAC TCC CAC CAC TG | TTG GAG TTG GAG TTG GAG TTG |
| ***Map1lc3a* (LC3B)** | GCC TGT CCT GGA TAA GAC CA | CCG TCT TCA TCC TTC TCC TG |
| ***Map1lc3b* (LC3A)** | CGT CCT GGA CAA GAC CAA GT | CAG GAA GCC GTC TTC ATC TC |
| ***Pik3c3*** | CTA ACG TGG AGG CAG ATG GT | CTG TCC AGC CAA TCC ACT TT |
| ***Tfeb*** | CTC AGT GGT CTT GGG CAA AT | TGT AGT CGA GGG GAG ACA GG |
| ***Ulk1*** | GCT CAC CTA AGC TGC CTG AC | ATT CTG AGA GCT GGG GGT TT |
| ***Arf4*** | CTG GCA AGA CGA CAA TTC TGT | CCA CAA AAA TGA GAC CCT GGG TA |
| ***Trappc13*** | CAA AGG ATG GCT CCA GGT TA | CAC GAG GTC CAT CAT CCT CT |



**Supplementary Table 3. PCR primer sequences used in the study (continued on the next page)**

| Primers | Forward (5' - 3') | Reverse (5' - 3') |
|---|---|---|
| ***Tnfrsf1b*** | ACA CCC TAC AAA CCG GAA CC | AGC CTT CCT GTC ATA GTA TTC CT |
| ***Tnfrsf10b*** | CGG GCA GAT CAC TAC ACC C | TGT TAC TGG AAC AAA GAC AGC C |
| ***Gorasp1*** | AGT CTG GGG TGT GGT ATT GG | TTG TGA GGT CGT AGC TGG AG |
| ***Gorasp2*** | GGG TTT ACA GAG GTC CAG CT | TGC CGA GCT AAT GGA GAG TC |
| ***Sqstm1*** | AGA ATG TGG GGG AGA GTG TG | TTT CTG GGG TAG TGG GTG TC |
| ***Gapdh*** | TGC ACC ACC AAC TGC TTA G | GGA TGC AGG GAT GAT GTT C |
| **Genotyping primers** | CTA CGG TGT AAA AGA GGC AGG | CTT GCG AAC CTC ATC ACT CGT |

**Supplementary Table 3. PCR primer sequences used in the study.**

| **Antibodies** | **Company** | **Catalogue #** | **Concentration**<br>**IB = immunoblot**<br>**IF = immunofluorescence** |
|---|---|---|---|
| α-smooth muscle actin | Abcam | ab5694 | IB(1:2000), IF(1:300) |
| α-tubulin | Santa Cruz Biotechnology | sc-5286 | IB(1:2000) |
| ATF4 | Cell Signaling Technology | 11815 | IB(1:1000) |
| β-actin | Cell Signaling Technology | 3700 | IB(1:4000) |
| Cre Recombinase | Cell Signaling Technology | 12830 | IB(1:200) |
| Desmin | Santa Cruz Biotechnology | sc-23879 | IF(1:200) |
| GAPDH | Millipore Sigma | MAB374 | IB(1:5000) |
| GFP | Abcam | ab6556 | IB(1:500), IF(1:300) |
| GFP (TRAP) | Bi-Institutional Antibody and Bioresource Core Facility | HtzGFP_02 (clone 19C8)<br>HtzGFP_04 (clone 19F7) | 50ug each per sample |
| Lamin A/C | Santa Cruz Biotechnology | sc-376248 | IB(1:2000), IF(1:300) |
| Lamin B1 | Santa Cruz Biotechnology | sc-30264 | IB(1:500) |
| LC3B | Cell Signaling Technology | 2775 | IB(1:1000) |
| MED6 | Santa Cruz Biotechnology | sc-390474 | IB(1:500) |
| MED25 | Santa Cruz Biotechnology | Sc-393759 | IB(1:500) |
| p62 | Cell Signaling Technology | 23214 | IB(1:1000) |
| PDGFRα | RnD Systems | AF1062 | IF(1:100) |
| PERK | Cell Signaling Technology | 3192 | IB(1:500) |
| phospho-eIF2α | Cell Signaling Technology | 3398 | IB(1:500) |
| eIF2α | Cell Signaling Technology | 5324 | IB(1:1000) |
| CHOP | Cell Signaling Technology | 2895 | IB(1:500) |
| sarcomeric actin | Invitrogen | MA1-21597 | IF(1:20) |
| Troponin T | Invitrogen | MA5-12960 | IF(1:200) |
| Vimentin | Cell Signaling Technology | 5741 | IF(1:300) |
| CREB3 | Proteintech | 11275-1-AP | IB(1:1000), IF(1:200) |
| RCAS1 | Proteintech | 66170-1-Ig | IF(1:200) |
| Donkey anti-goat 594 | Invitrogen | A-11058 | IF(1:400) |
| Goat anti-mouse 594 | Invitrogen | A-21044 | IF(1:400) |
| Goat anti-rabbit 488 | Invitrogen | A-11034 | IF(1:400) |
| Goat anti-rabbit 594 | Invitrogen | R-37117 | IF(1:8) |
| Licor green mouse | LI-COR Biosciences | 926-32210 | IB(1:5000) |
| Licor green rabbit | LI-COR Biosciences | 926-32211 | IB(1:5000) |
| Licor red mouse | LI-COR Biosciences | 926-68070 | IB(1:5000) |
| Licor red rabbit | LI-COR Biosciences | 926-68071 | IB(1:5000) |



**Supplementary Table 4. Antibodies and their dilutions used in the study.**
